# The cell adhesion molecule Sdk1 shapes assembly of a retinal circuit that detects localized edges

**DOI:** 10.1101/2021.06.08.447607

**Authors:** Pierre-Luc Rochon, Catherine Theriault, Aline Giselle Rangel Olguin, Arjun Krishnaswamy

## Abstract

Nearly 50 different mouse retinal ganglion cell (RGC) types sample the visual scene for distinct features. RGC feature selectivity arises from its synapses with a specific subset of amacrine (AC) and bipolar cell (BC) types, but how RGC dendrites arborize and collect input from these specific subsets remains poorly understood. Here we examine the hypothesis that RGCs employ molecular recognition systems to meet this challenge. By combining calcium imaging and type-specific histological stains we define a family of circuits that express the recognition molecule Sidekick 1 (Sdk1) which include a novel RGC type (S1-RGC) that responds to local edges. Genetic and physiological studies revealed that Sdk1 loss selectively disrupts S1-RGC visual responses which result from a loss of excitatory and inhibitory inputs and selective dendritic deficits on this neuron. We conclude that Sdk1 shapes dendrite growth and wiring to help S1-RGCs become feature selective.

## Introduction

In the retina, each of the ∼46 types of retinal ganglion cell (RGC) synapses with a unique subset of amacrine (AC) and bipolar cell (BC) types to create circuits that detect a unique aspect of the visual scene (Clark and Demb, 2016; Gollisch and Meister, 2010; Jadzinsky and Baccus, 2013; Sanes and Masland, 2015; Shekhar et al., 2016; Tran et al., 2019; Yan et al., 2020). A growing body of work suggest that recognition molecules guide the neurites of newborn retinal neurons to grow into sublayers of a specialized neuropil, the inner plexiform layer (IPL), where they contact hundreds of potential synaptic targets (Rangel Olguin et al., 2020; Sanes and Yamagata, 2009; Sanes and Zipursky, 2020; Zhang et al., 2017).

The factors that guide developing arbors to synapse with appropriate targets within a layer are not well understood. An initial idea, called Peter’s Principle, posited that developing neurons synapse with nearby cells according to how often they make contact (Binzegger et al., 2004; Peters and Feldman, 1976; Shepherd et al., 2005). However, recent connectomic studies of the IPL demonstrate no obvious relationship between contact frequency and synapse number (Briggman et al., 2011; Helmstaedter et al., 2013). Instead, these connectomic data support a model in which neurons recognize targets in their immediate vicinity and synapse specifically with them. Key molecules in this recognition process are thought to be members of the immunoglobulin superfamily (IgSFs).

Briefly, IgSFs are adhesion molecules that bind to themselves (homophillic) or compatible IgSFs (heterophilic) across cell-cell junctions. It has been proposed that selective IgSF expression within synaptic partners allows their neurites to adhere and synapse when they encounter each other (Sanes and Zipursky, 2020). A recent study in mouse retina provides support for this view (Krishnaswamy et al., 2015). In this study, the IgSF Sidekick-2 (Sdk2) in VG3-ACs and W3B RGCs, drives these neurons to synapse with each other far more than they synapse with the Sdk2 negative neurons they contact. Loss of Sdk2 ablates the enhanced VG3-W3B connectivity but does not alter the gross structure or overlap of their arbors, suggesting that IgSFs increase the probability of synapses between this pair (Krishnaswamy et al., 2015). On the other hand, direction selective RGCs require the IgSF Contactin 5 (Cntn5) to grow dendritic branches in IPL layers bearing axons of their AC/BC partners. Loss of Cntn5 from these RGCs decreases dendritic branches and reduces synaptic input (Peng et al., 2017), suggesting that IgSFs influence connectivity by regulating intra-laminar dendritic growth. These studies suggest a common role for these IgSFs to stabilize/promote the growth of small dendrites that lead to synapses or suggest differing roles for IgSFs in synaptic specificity and neurite growth. Too few IgSFs have been studied in the context of mammalian circuit assembly to draw a firm conclusion. To gain more insight, we investigated the closest IgSF relative of Cntn5 and Sdk2, called Sidekick-1 (Sdk1) in retinal circuit assembly.

We show that Sdk1 is expressed by a family of 5 RGCs whose dendrites target IPL layers bearing the processes of 5 Sdk1-expressing interneurons (ACs and BCs). We uncover molecular markers for each Sdk1 RGC and applied these markers following calcium imaging to investigate their visual responses, discovering that the Sdk1 family includes two ON-direction selective RGCs and an RGC (S1-RGC) sensitive to local edges. Loss of Sdk1 disrupted responses to visual stimuli on S1-RGCs without affecting other members of the Sdk1-RGC family. Using electrophysiological recordings, we show that Sdk1 loss reduced excitatory and inhibitory synaptic inputs to S1-RGCs, decreasing its firing to visual stimuli. These synaptic deficits were specific to S1-RGCs as stimulus evoked responses on Sdk1+ ON-alpha RGCs were unaffected by Sdk1 loss. Finally, we show that the loss of Sdk1 does not alter the IPL-layer targeting of S1-RGC dendrites but selectively reduces their intralaminar complexity and size. From our results, we conclude that Sdk1 is required for S1-RGC to develop dendritic arbors and receive AC and BC synapses.

## Results

### Sdk1 labels a family of retinal circuits

To investigate the expression of Sdk1 across the retina, we used mice in which the Sdk1 gene is disrupted by the presence of cDNA encoding a Cre-GFP fusion protein (Sdk1^CG^); as heterozygotes these lines allow selective access to Sdk1 neurons, as homozygotes they are Sdk1 nulls (Krishnaswamy et al., 2015; Yamagata and Sanes, 2018). Cross sections and wholemounts from these animals showed GFP+ nuclei within subsets of neurons expressing the BC marker Chx10, the AC marker Ap2-α, and the RGC marker RBPMS (Figure 1A-F); no labeling was found in horizontal cells and photoreceptors. Counting double-labelled cells showed ∼7795 Sdk1-BCs//mm^2^, ∼350 Sdk1-ACs /mm^2^, and ∼390/mm^2^ Sdk1-RGCs (Figure 1G), suggesting that Sdk1 labels several retinal circuits. To address this, we set out to match GFP+ neurons to retinal subtypes.

**Figure 1.**
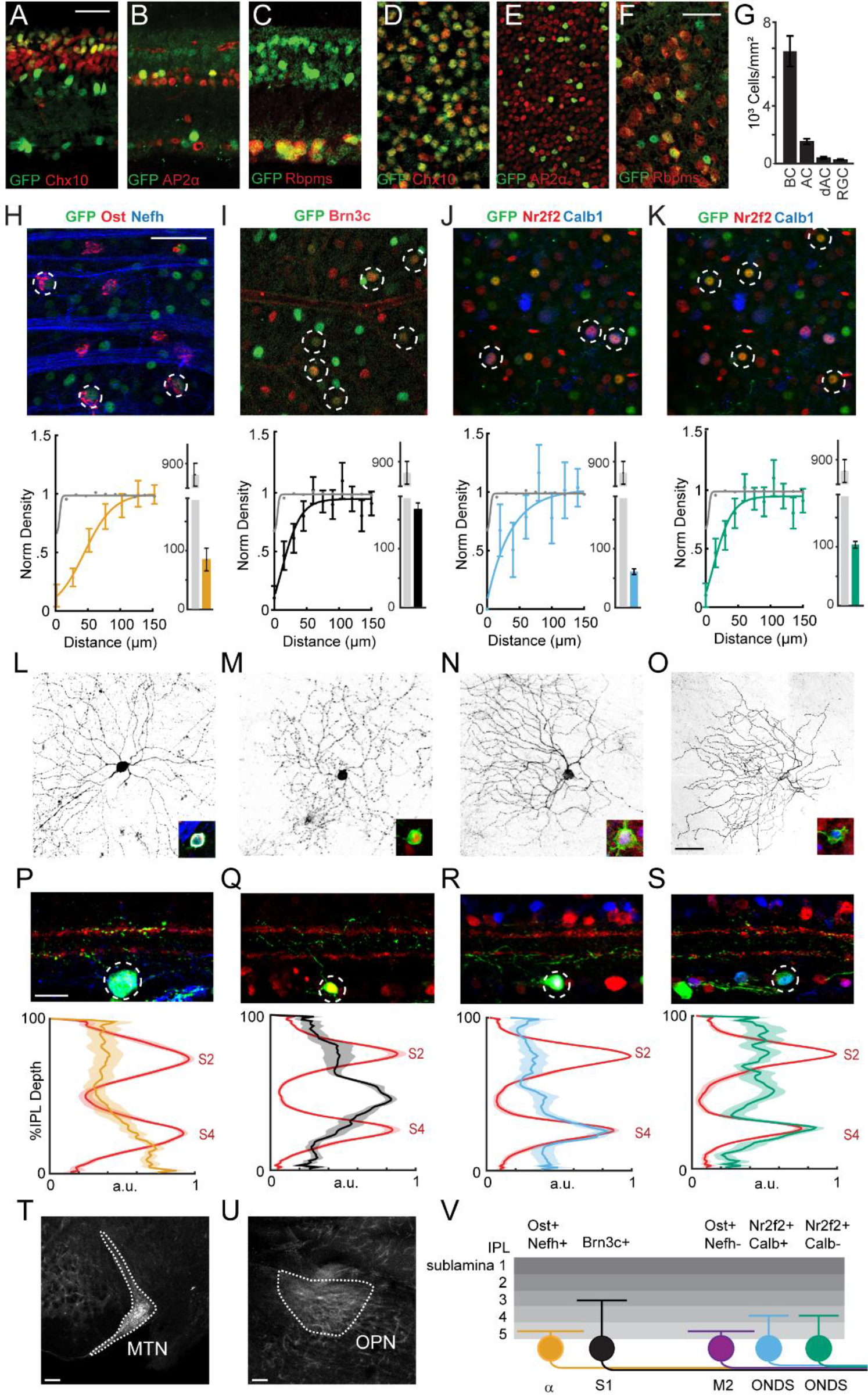
Sdk1 labels a family of RGC types. **A-F.** Sample cross-sections (A-C) and wholemounts (D-F) from Sdk1^CG^ mice stained with antibodies to GFP and the BC marker Chx10 (A, D), AC marker AP2 (B,E), and RGC marker RBPMS (C,F). Scale = 25μm. **G.** Bar graph showing density of BCs, ACs, displaced amacrine cells (dACs), and RGCs expressing Sdk1 computed from experiments like those shown in D-F. (n= 14 fields from 3 animals for each experiment) **H-K.** Top: GFP-stained sample Sdk1^CG^ retinal wholemounts co-stained with antibodies to osteopontin (Ost) and neurofilament heavy chain (Nefh) (H), Brn3c (I), or Nr2f2 and calbindin (Calb) (J-K). Dotted circles indicate co-labelled neurons and Scale = 50μm. Bottom: normalized density recovery profiles and average density of co-labelled RGCs measured from corresponding experiments shown in the top row. Colored traces indicate density recovery profiles for co-labeled neurons; gray traces indicated density recovery profiles for all GFP+ cells in the GCL. (n= 13-18 fields from 6 animals for each experiment) **L-O.** Images showing the dendritic morphology of individually labelled Ost+/Nefh+ RGCs (L), Brn3c+ RGCs (M), Nr2f2+/Calb+ RGCs (N), and Nr2f2+/Calb− RGCs (O) in Sdk1^CG/+^ retinal wholemounts. Inset shows marker expression in the soma. **P-S.** Top: Images showing the laminar morphology of individually labelled Ost+/Nefh+ RGCs (P), Brn3c+ RGCs (Q), Nr2f2+/Calb+ RGCs (R), and Nr2f2+/Calb− RGCs (S) in Sdk1^CG/+^ retinal cross sections. Red staining indicates VAcht, a marker for sublaminae 2 (S2) and 4 (S4) (Scale = 50μm). Bottom: IPL linescans measured from corresponding experiments shown in the top row. Red traces show VAchT intensity and colored traces show reporter intensity measured from experiments like those shown in the top row. (n = 6-12 fields per RGC type from more than 16 animals) **T-U.** Magnified brain images taken from Sdk1^CG^ mice intraocularly injected with Cre-dependent reporters showing Sdk1-RGC innervation in the medial terminal nucleus (MTN) (T) and olivary pretectal nuclei (OPN) (U). Scale = 100μm. **V.** Summary cartoon of the Sdk1 RGC family showing Ost+/Nefh+ Onα-RGCs, Brn3c+ S1-RGCs, Ost+/Nefh− M2-RGCs, and Nr2f2+/Calb+ and Calb- ON-DSGCs.

#### Sdk1 defines a family of 5 RGCs

Publicly available single RGC sequencing atlases indicate that Sdk1 is highly expressed in ∼6 RGC clusters (Rheaume et al., 2018; Tran et al., 2019). Two clusters correspond to alpha RGCs with sustained responses to light onset (ONα-RGCs) and type-2 melanopsin-positive RGCs (M2-RGCs). Four others predicted novel RGCs types (Figure 1 – figure supplement 1A). To map these clusters onto GFP+ RGCs, we compared Sdk1 clusters to all RGCs and identified 5 genes that combinatorially label each Sdk1-RGC type: (1) Onα-RGCs should strongly express neurofilament heavy chain isoform (Nefh), osteopontin (Ost) and the intracellular calcium buffer calbindin (Calb); (2) M2-RGCs should express high levels of Ost+, low levels of Nefh, and low levels of Calb; (3) two predicted novel RGCs should express the steroid hormone receptor (Nr2f2), (4) with one of these also expressing Calb; (5) another novel RGC should express the Pou-domain containing transcription factor (Brn3c); and (6) a final RGC should not express any of these genes (Figure 1– figure supplement 1B).

To validate these predictions, we took advantage of the mosaic arrangement of retinal types. Briefly, retinal neurons of the same type are spaced apart at a characteristic distance, whereas neurons of different types are spaced randomly. Thus, the density of cells labelled with a candidate marker will drop at short distances from a reference cell if the candidate labels a single type. Sdk1^CG^ wholemounts stained with antibodies to Nefh and Ost showed a pair of Nefh+/Ost+ and Nefh-/Ost+ mosaics spaced at distances expected for the ONα-RGC (Figure 1H) and M2-RGC (Figure 1 –figure supplement 1C), respectively. A high-density mosaic was labelled by Brn3c (Figure 1I), and a final pair of mosaics was found to be Nr2f2+/Calb− and Nr2f2+/Calb1+ (Figure 1J-K). We were unable to find an RGC that corresponded to the sixth Sdk1 cluster, as a stain with all these RGC markers and Ap2-α labelled all GFP+ neurons in the GCL (Figure 1 – figure supplement 1E-G). Thus, we provide molecular definition for 5 Sdk1-RGCs, which include 3 predicted novel types.

We used two approaches to define the anatomy of Sdk1- RGCs. In one approach, retrogradely infecting AAVs bearing Cre-dependent reporters were delivered to the lateral geniculate nucleus (LGN) or superior colliculus (SC) of Sdk1^CG^ mice. In the other, tamoxifen was used to drive reporter expression in a related strain (Sdk1^CE^) whose Sdk1 gene is disrupted with cDNA encoding a Cre-human estrogen receptor (CreER) fusion protein (Krishnaswamy et al., 2015). Staining retinae from these mice with our panel of molecular markers revealed the morphology of two Sdk1-RGCs: (1) Ost+/Nefh+ RGCs bore large somas and wide dendritic arbors confined to IPL sublamina 5 and matched previous descriptions of ONα-RGCs (Figure 1L,P) (Krieger et al., 2017; Krishnaswamy et al., 2015); and (2) Brn3c+ RGCs had small somas and grew dendritic arbors confined to the center of sublamina 3 (Figure 1M,Q). We refer to these Brn3c+ neurons S1-RGCs. Ost+/Nefh- and Nr2f2+ RGCs were rarely observed using this method. These results indicate that only a subset of Sdk1-RGCs project strongly to the LGN and SC.

One possibility for the rare observation of Ost+/Nefh- and Nr2f2+ RGCs is that these RGCs innervate ‘non-imaging forming’ brain regions more strongly than they innervate the LGN and SC (Martersteck et al., 2017). Consistent with this, brain sections taken from intraocularly infected Sdk1^CG^ mice showed reporter-labelled RGC axons in the non-image forming medial terminal nucleus (MTN) and olivary pretectal nuclei (OPN) (Figure 1T-U). To sparsely label these RGCs, we tamoxifen treated Sdk1^CE/+^ mice crossed to Cre-dependent reporters and stained their retinae with Ost, Nefh, Nr2f2 and Calb. These experiments revealed that both Nr2f2+/Calb+ and Nr2f2+/Calb- RGCs grew oblong dendritic arbors confined to IPL sublamina 4 (Figure 1N, R, O, S). We obsevered a few Ost+/Nefh- RGC somas in fields labelled densely by reporter (Figure 1 –figure supplement 1H), but we were unable to mark them individually, likely due to their low density (∼20/mm^2^) (Sanes and Masland, 2015). Thus, Sdk1 labels a family of 5 RGCs which include image- and non-image forming types.

#### Sdk1 defines at least 5 interneuron types

We next asked if Sdk1-ACs and BCs target the same layers as Sdk1-RGCs. Staining reporter-labelled retinae from Sdk1^CE/+^ mice with Ap2-α revealed 3 kinds of Sdk1-ACs (Figure 1 –figure supplement 2A-C). One of these had very large (>1mm) diameter dendritic arbors located in sublamina 5 (Figure 1 –figure supplement 2A), another had a similarly large arbor located in sublamina 3 and resembled descriptions of type 2 catacholaminergic ACs (Figure 1 –figure supplement 2B) (Knop et al., 2014; Krishnaswamy et al., 2015). A third Sdk1-AC grew comparatively narrower dendritic arbors that exhibited the characteristic waterfall appearance of the A17 AC (Figure 1 –figure supplement 2C) (Masland, 2012). Cross-sections from the same experiments also showed the presence of many rod BCs as judged by their anatomy (Figure 1 – figure supplement2D), as well as many BCs that targeted the interface between sublamina 3 and 4, matching reports of Type 7 BCs (Figure 1 –figure supplement2E) (Wassle et al., 2009). The IPL lamination profiles of Sdk1-ACs, BCs and RGCs show that these cells target a common set of IPL laminae. Thus, we conclude that Sdk1 defines a family of neurons whose circuits are contained within the inner 3 layers of the IPL.

### Sdk1-RGCs include direction selective and edges-detecting subtypes

#### Registering calcium imaged fields to posthoc stains assigns molecular identity to RGC response

To characterize the functional properties of the Sdk1 RGC family, we devised a procedure to relate marker-gene expression to neural response (Figure 2A, Figure 2 –figure supplement 1). Briefly, retinal neurons in Sdk1^CG^ mice were intraocularly infected with AAVs bearing Cre-dependent genetically encoded calcium indicators (GCaMP6f) and two weeks later, we imaged their responses to full-field flashes and bars moving at high or low-velocity in 8 different directions using two-photon microscopy. To register each two-photon field to the optic nerve head, we labeled blood vessels fluorescently labelled with sulphorhodamine101 and matched the vascular pattern (Figure 2 –figure supplement 1A-B). Next, we fixed and stained retinae for GFP, Sdk1-RGC markers, and vessels, and re-imaged these retinae with a confocal microscope (Figure 2 –figure supplement 1C). Vascular patterns let us register confocal and two-photon fields (Figure 2 – figure supplement 1F-K) and Ost distribution let us orient each retina along the dorso-ventral and naso-temporal axes (Bleckert et al., 2014). Finally, ROIs were drawn using the confocal image and applied to channels containing marker stains and to the two-photon images to extract responses (Figure 2 –figure supplement 1L-O)

**Figure 2.**
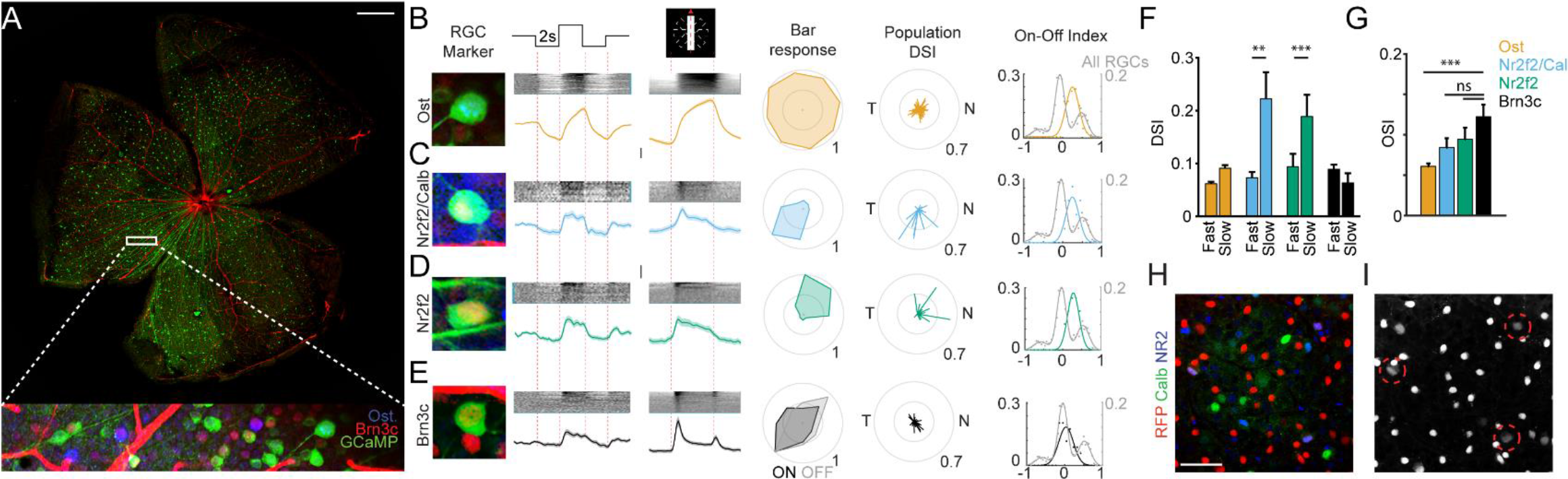
Function of Sdk1 circuits. **A.** Sample wholemount from a Sdk1^CG^ retina infected with AAV-GCaMP6f labelled with the positions of a typical two-photon field (Scale = 500μm). Inset magnifies the boxed field after immunostaining and shows GCaMP labelled Sdk1-RGCs (green) stained with Brn3c (red) and Osteopontin (Ost, blue). **B-E.** Magnified somata image, average full-field response, average bar response, sample polar plot, population direction selective indices (DSI) and on-off index for Ost+ (B, n = 107 cells from 18 retinae), Nr2f2+/Calb+ (C, n = 41 cells from 6 retinae), Nr2f2+/Calb− (D, n = 34 cells from 6 retinae), and Brn3c+ RGCs (E, n = 95 cells from 6 retinae). Each RGC shows a characteristic pattern of responses to stimuli. On-Off index distribution from a mouse line in which all RGCs express GCaMP6f (All RGCs; n = 1426cells from 5 retinae) is shown for comparison. Vertical scales = z-score of 2. **F.** Average DSI computed from bar stimuli moving at ∼1000mm/sec (fast) or ∼200mm/sec (slow) for each Sdk1-RGC group. Significance: ** = p<0.01; *** = p<0.001. **G.** Average orientation selective indices computed from bar stimuli for each Sdk1-RGC group. **H-I.** Wholemount retina from the Pcdh9-Cre line crossed to Cre-dependent reporter stained for Nr2f2 and Calb (E). Arrows show triple labelled cells (F). (Scale = 50μm)

A balance between antibody species restrictions and throughput led us to group the low-abundance M2-RGC (<20/mm^2^) with ONα-RGCs using the marker Ost. This registration procedure divided our GCaMP6f dataset into 4 marker-defined groups, with each bearing a characteristic response to visual stimuli and enrichment of RGC markers (Figure 2, Figure 2 – figure supplement 2A-D). To check the consistency of these marker defined groups, we compared the bar-evoked responses within each group to the average response of all four groups and asked if grouped traces were most similar to their group mean. To do this, we normalized each trace and computed its cosine similarity to the mean response of each group. Ost+ and Brn3c+ traces showed highest similarity to their own mean group responses. Nr2f2+ and Nr2f2+/Calb+ traces also showed high similarity to their mean group response but also showed similarity to each other’s mean response, consistent with the strong resemblence in their visual responses (Figure 2 –figure supplement 2A-D). These results indicate that our registration procedure can group molecularly and functionally similar RGCs.

#### Ost+ RGCs show sustained responses to light onset

Ost+ RGCs (M2- and Onα -RGCs) showed sustained responses to the onset of a full-field flash and leading edge of the moving bar. Converting these bar responses to an ON-OFF index (see methods) and comparing the distribution of these indices to those computed from a dataset in which all RGCs express GCaMP6f (Vglut2-Cre/LSL-GCaMP6f) emphasize this observation (Figure 2B). Plotting bar direction versus evoked response on a polar plot showed no obvious preference for motion direction (Figure 2B). Converting each polar plot to a direction selective index (DSI) and preferred angle and viewing the entire Ost+ population on polar axes confirmed this picture, showing weak directional tuning with no systematic preference for stimuli moving along any cardinal axis (Figure 2B). Comparing the average DSI within this group for fast- or slow-moving bars was similar, indicating a lack of motion selectivity within the Ost+ RGC group (Figure 2F). These results are consistent with previous reports of ONα- and M2-RGCs (Berson et al., 2010; Krieger et al., 2017).

#### Nr2+ RGCs are ON-direction selective

On the other hand, both Nr2f2+ RGC types gave transient responses to the leading edge of the moving bar that varied systematically with bar direction. Aligning these responses to retinal orientation revealed polar plots that pointed ventrally for Nr2+/Calb+ RGCs (Figure 2C) or moving dorsally for Nr2+/Calb- RGCs (Figure 2D). DSI polar plots for these two populations showed the same respective biases for ventral or dorsal motion and reducing bar velocity caused the average DSI of both RGC types to increase (Figure 2F), indicating that Nr2f2+ RGCs encode the direction of slow-moving bright stimuli. Taken together with our anatomical results, Nr2f2+ RGCs strongly resemble ON-direction selective RGCs (ON-DSGCs) which grow dendrites in sublamina 4 and comprise 3 subtypes attuned to motion towards either dorsal, ventral, and nasal poles of the retina (Dhande et al., 2013; Sanes and Masland, 2015; Yonehara et al., 2009). Staining reporter-labelled retinae from the Pcdh9-Cre line, which marks ventrally-tuned ON-DSGCs (Lilley et al., 2019; Matsumoto and Yonehara, 2018) showed overlap with ventral-motion selective Nr2f2+/Calb+ RGCs (Figure 2H-I). Thus, we conclude that Nr2f2+ RGCs likely correspond to a pair of ON-DSGCs.

#### Brn3c+ S1-RGCs are ON-OFF RGCs that respond to local edges

S1-RGCs (Brn3c+) showed ON responses to a full field flash but responded to both the leading and trailing edge of the bright moving bar (Figure 2E). These results indicate that S1-RGCs can respond to both bright and dark stimuli that are localized to their receptive field. Larger stimuli appear to recruit surround mechanisms, which strongly attenuate S1-RGCs, particularly its OFF responses. Plotting S1-RGC bar responses against bar direction showed that many cells respond to bars traveling along the same axis, suggesting that these neurons detect stimulus orientation (Figure 2E). Computing an orientation selectivity index (OSI) for these cells showed a higher orientational preference in this neuron as compared with the other Sdk1-RGCs (Figure 2I), however, these values are significantly weaker than the OSI found in recently described orientation-selective RGCs (Nath and Schwartz, 2016, 2017). Thus, we conclude that S1-RGCs are ON-OFF cells that respond best to edges that fall within their receptive fields.

### Sdk1 loss selectively impairs S1-RGCs visual responses

#### Sdk1 loss impairs a subset of types in the Sdk1 RGC family

IgSFs like Sdk1 are thought to guide RGCs to synapse with specific AC and BC types to create feature-selective neural circuits (Sanes and Zipursky, 2020). Given that Sdk1 labels a family of 5 RGCs and interneurons whose processes overlap in a common set of lamina, we next asked if Sdk1 is important for the development of functional properties in Sdk1 RGCs. To test this idea, we labelled Sdk1 neurons with GCaMP using intraocular AAV injections in Sdk1^CG/CG^ mice, imaged their responses, and grouped these responses using type-specific markers. The responses of Sdk1-null Ost+ RGCs, which include Onα- and M2-RGCs, resembled those of their control counterparts (Figure 3A). Nr2f2+ RGCs, which include two kinds of ON-DSGCs, showed subtle changes in their average responses to full-field and moving bar stimuli (Figure 3B-C) with somewhat altered preference and selectivity for motion in the NR2f2+/Calb+ group (Figure 3B). S1-RGCs were most affected by Sdk1 loss and showed a significant decrease in response magnitude to bar stimuli (Figure 3D). These results indicate that Sdk1 loss impairs S1-RGCs but leaves the other Sdk1-RGCs relatively unaffected. Interestingly, single-cell sequencing data suggested that the other Sdk1-RGCs express high levels of Sdk2, suggesting that there could be functional redundancy in Onα-, M2-, and ON-DSGCs (Figure 3 –figure supplement 1A). We partially confirmed this expression pattern by staining retinae from a Sdk2-CreER knock-in line crossed to reporters with Ost, Nefh, Nr2f2 and Calb antibodies and found many Onα-, and Nr2f2+/Calb+ RGCs (Figure 3 –figure supplement 1B-C). These results suggest that the higher levels of Sdk2 in Onα, M2 and ON-DSGCs compensates for Sdk1 loss and attenuates phenotypes. Based on these results, we focused on the S1-RGC for the rest of this study.

**Figure 3.**
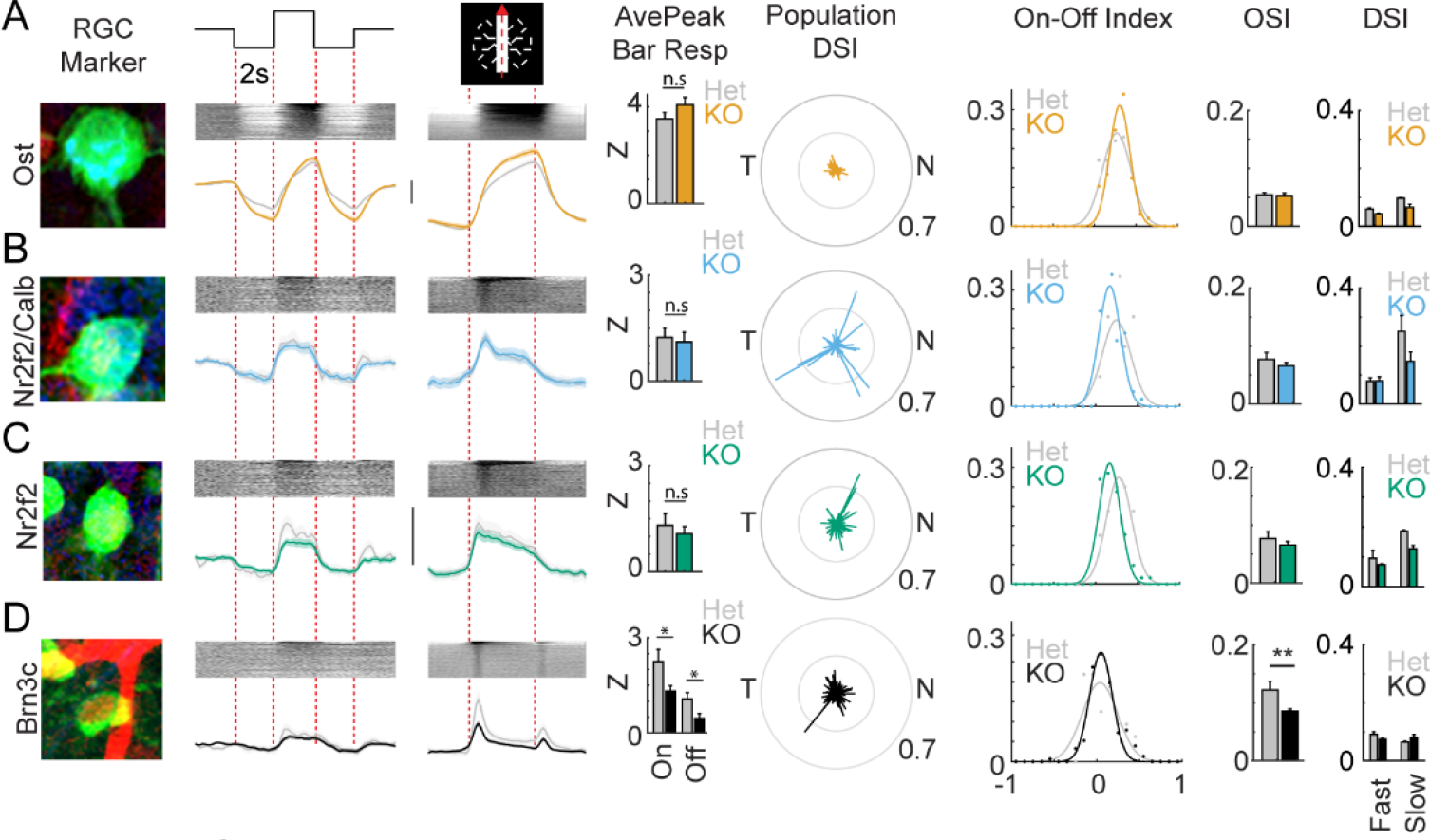
Sdk1 loss causes selective deficits in S1-RGCs. **A-D.** Magnified somata image, average full-field response, average bar response, average peak response to bars, population direction selective indices (DSI), on-off index, mean orientation selective index (OSI) and mean direction selective index (DSI) corresponding to Ost+ (A, n = 97 cells from 10 retinae), Nr2f2+/Calb+ (B, n = 59 cells from 6 retinae), Nr2f2+/Calb− (C, n = 77 cells from 6 retinae), and Brn3c+ RGCs (D, n = 147 cells from 7 retinae) in Sdk1^CG/CG^ retinae. Grayed traces and bars show the same measurements from Sdk1 heterozygotes (Het) (Vertical scales = z-score of 2). Sdk1 loss alters Brn3c RGC visual responses. Significance: * = p<0.05.

#### Sdk1 loss selectively disrupts S1-RGC visual responses

Retrograde viral injections in the LGN/SC of Sdk1^CG/+^ offered us a way to study the role of Sdk1 in S1-RGCs and use Sdk1+/Sdk2+ ONα-RGCs as an internal comparison (Figure 4A-B). Recordings from Sdk1^CG/+^ mice showed many RGCs whose anatomical and functional properties matched one of these two groups. S1-RGCs were easily targeted for loose-patch recordings by their small soma and showed ON and OFF responses to a moving bar that were strongest for motion along a single axis as judged by their elongated profiles on a polar plot (Figure 4A-C). Similar recordings from nearby large-soma RGCs showed sustained responses to light onset with little tuning for motion direction (Figure 4C).

**Figure 4.**
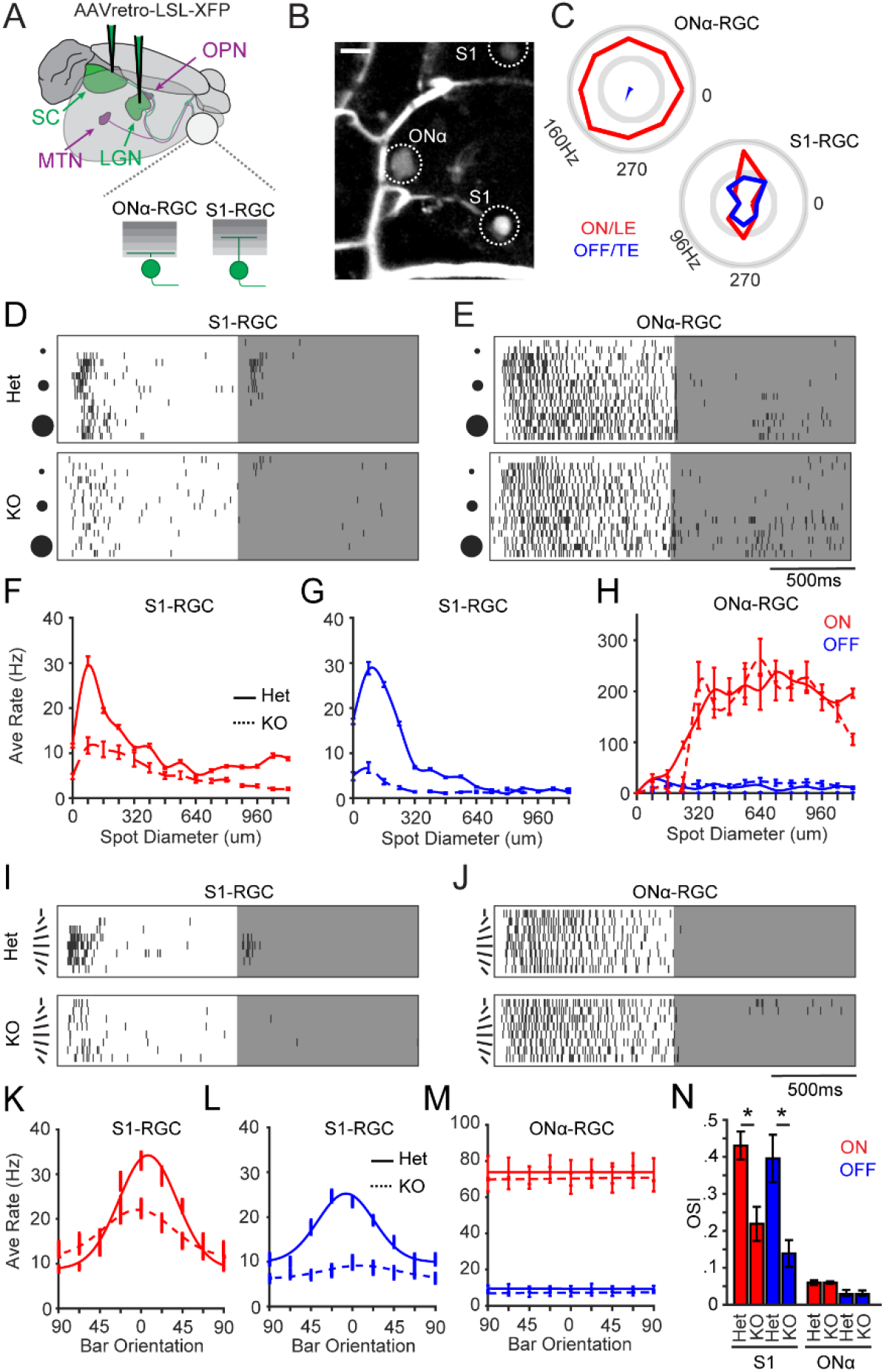
Selective loss of visual responses on S1-RGCs in the absence of Sdk1. **A.** Cartoon of S1-RGCs and ONα-RGCs labelled by delivering retrogradely infecting AAVs bearing cre-dependent reporters in the lateral geniculate nucleus (LGN) or superior colliculus (SC) of Sdk1^CG^ mice. Targets of other Sdk1-RGCs that project to olivary pretectal nuclei (OPN) and medial terminal nucleus (MTN) are also shown. **B.** Sample two-photon image of a Sdk1^CG/+^ retina labelled as described in A showing a large soma ONα-RGC and a small soma S1-RGC. Sulphorhodamine101 labels vessels in the GCL. Scale = 25μm. **C.** Polar plots of spike responses to a bar moving in 8 different directions recorded from example ONα- and S1-RGCs in experiments like those shown in B. **D-E.** Raster of spike responses to an expanding flashing spot recorded from example S1-RGCs (D) and Onα-RGCs (E) in Sdk1^CG/+^ (Het) or Sdk1^CG/CG^ retinae (KO). **F-G.** Average S1-RGCs firing rates versus bright (F) or dark (G) spot diameter measured from experiments like those shown in D. Average Onα-RGCs firing rate versus bright (ON) or dark (OFF) spot diameter measured from experiments like those shown in E. **I-J.** Raster of spike responses to centered dark or bright bar rotating through 8 orientations recorded from S1-RGCs (I) and Onα-RGCs (J) in Sdk1^CG/+^ (Het) or Sdk1^CG/CG^ retinae (KO). **K-L.** Average firing rate versus bar orientation for S1-RGCs measured from experiments like those shown in I. **M.** Average firing rate versus bar orientation for ONα-RGCs measured from experiments like those shown in J. **N.** Average orientation selectivity indices computed for S1-RGC and ONα- RGC responses to the oriented bar stimulus in control (Het) and Sdk1-null (KO) retinae (n=7 for Sdk1^CG/+^ ON RGCs, n=12 for Sdk1^CG/+^ Brn3c RGCs, n=14 for Sdk1^CG/CG^ Brn3c RGCs, n=6 for Sdk1^CG/CG^ ONα RGCs; * = p <0.05).

Our calcium imaging experiments showed that S1-RGCs have strong surround suppression. To relate these signals to spiking behavior, we recorded responses from control S1-RGCs to an expanding spot centered over their receptive field that flashed on and off. S1-RGCs showed ON and OFF responses to this stimulus that were strongly suppressed by spot size, with OFF responses fully suppressed by spots exceeding ∼300μm (Figure 4D, F-G). Both ON and OFF components were most strongly activated by spot sizes close to the diameter of S1-RGC dendritic arbors, indicating that the receptive field center on these neurons is ∼120-150μm (Figure 4F-G). Recordings from control ONα-RGCs displayed higher baseline spiking than S1-RGCs and sustained ON responses that were poorly suppressed by large stimuli (Figure 4E), consistent with their full-field and moving bar responses in our calcium imaging experiments. In contrast, the same recordings from S1-RGCs in Sdk1^CG/CG^ retinae showed a dramatic loss of responsivity to dark stimuli and significantly weaker responses to ON stimuli (Figure 4D, F-G). Recordings from nearby Sdk1-null ONα-RGCs showed comparable responses to their control counterparts (Figure 4E, H), indicating that Sdk1 loss selectively affects S1-RGCs.

We found that most S1-RGCs showed responses to axial bar motion, similar to what we saw in our calcium imaging experiments. We suspected the inability to align bars with S1-RGC receptive fields in the imaging studies could have activated their strong surround and attenuated their responses to this stimulus. We revisited the idea that these neurons might exhibit sensitivity to stimulus orientation, presenting stationary bars whose width matched the receptive field size of S1-RGCs and rotated through 8 different orientations. As expected from their responses to moving bars, S1-RGCs responded preferentially to both bright and dark bars oriented along a single axis (Figure 4I) whereas Onα-RGCs respond to bright bars alone with little response variation to bar orientation (Figure 4J). The same stimulus evoked poor responses from S1-RGCs in Sdk1^CG/CG^ retinae (Figure 4I, K-L), with significantly weaker responses to dark and bright bars, confirming our results using bright and dark expanding spot stimuli. The same recordings from nearby Sdk1-null Onα- RGCs showed similar responses to those in controls (Figure 4J, M). Computing OSI values for these cells and comparing the mean OSI for S1-RGCs and ONα-RGCs in controls and knockout retina showed a selective reduction of orientation selectivity for S1-RGCs in the absence of Sdk1 (Figure 4N). Thus, we conclude that Sdk1 is required for S1-RGCs to develop normal responses to visual stimuli.

#### Sdk1 loss impairs excitatory and inhibitory synaptic inputs to S1-RGCs

The deficits we observed on S1-RGCs in Sdk1^CG/CG^ retinae might result from a loss of excitatory inputs, a change in inhibitory inputs, or both. To investigate this idea, we recorded synaptic currents from S1-RGCs and ONα-RGCs in control and Sdk1 null retinae and compared their responses to visual stimuli. We began with expanding spots, isolating excitatory and inhibitory inputs by holding neurons at −60mV and 0mV respectively (Figure5A-B). S1-RGCs in controls showed inward (excitatory) currents at −60mV to both bright and dark spots that grew in strength until the spot reached ∼150μm in diameter and then steadily weakened (Figure 5C-D). Outward (inhibitory) currents at 0mV were also found to both bright and dark spots but grew steadily with spot diameter (Figure 5E-F). Recordings from S1-RGCs in Sdk^CG/CG^ retinae showed significantly weaker inward and outward currents to bright and dark spots at all spot diameters (Figure 5A-F), indicating a loss of functional excitatory and inhibitory synapses on S1-RGCs in the absence of Sdk1. Whole-cell recordings from nearby ONα-RGCs showed inward and outward currents that were similar between controls and knockouts (Figure 5 – figure supplement 1A-B), indicating that Sdk1-loss impairs synaptic inputs on S1-RGCs rather than generally affecting excitatory and inhibitory synaptic strength.

**Figure 5.**
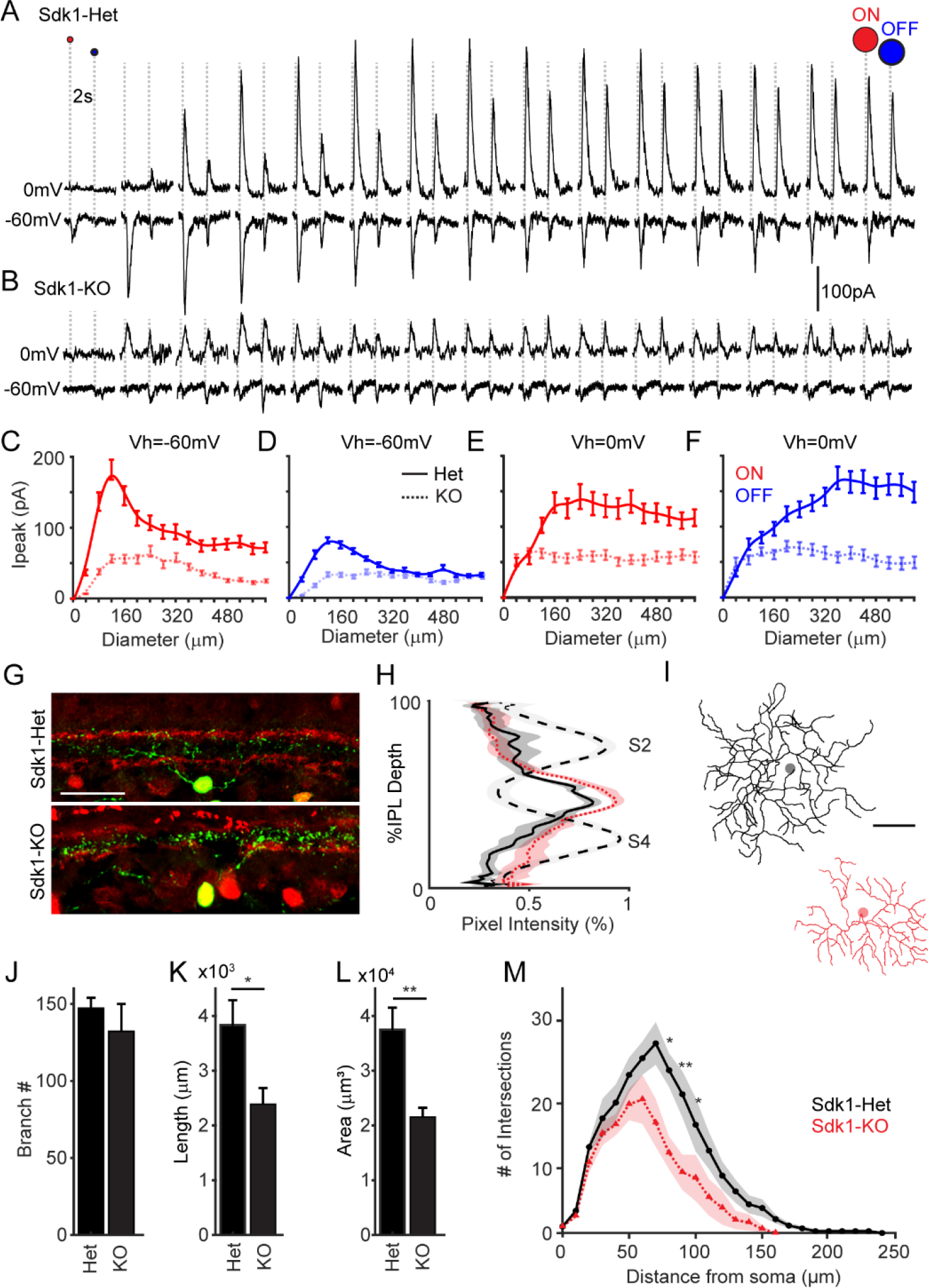
Sdk1 loss causes S1-RGCs to lose synaptic inputs and dendritic arbor complexity. **A-B.** Whole-cell recordings from S1-RGCs held at potentials to isolate excitatory (∼ −60mV) or inhibitory (∼0mV) currents to an expanding flashing spot in Sdk1^CG/+^ (Het) or Sdk1^CG/CG^ retinae (KO). **C-D.** Average peak current versus expanding bright (C) or dark (D) spot diameter measured from control or knockout S1-RGCs held at −60mV in experiments like those shown in A-B. **E-F.** Average peak current versus expanding bright (E) or dark (F) spot diameter measured from control or knockout S1-RGCs held at −0mV taken experiments like those shown in A-B. (n=8 for Sdk1-Het, n=9 for Sdk1-KO) **G.** Retinal cross-sections showing S1-RGCs in control (Het) Sdk1 knockout (KO) retinae. Scale = 50μm. **H.** Linescans through S1-RGC arbors in control in Sdk1 null retinae taken from experiments like those shown in G. **I.** Skeletonized S1-RGC dendrites from control (black) and Sdk1 null (red) retinae. (Scale = 50μm). **J-L.** Average branch number (J), branch length (K), and dendritic area (L) measured from control and Sdk1 null S1-RGC dendritic arbors. **M.** Sholl analysis of dendritic arbors measured from Het and KO S1-RGCs. (n = 8 for both Sdk1-Het and Sdk1-KO; * = p < 0.05; ** = p < 0.01)

Finally, we examined the synaptic currents evoked on S1-RGCs to our oriented bar stimuli. Controls showed inward and outward currents to both bright and dark oriented bars, but the magnitudes of these currents varied with bar orientation (Figure 5 –figure supplement 2A-B). Integrating these currents across the presentation time of each bar and normalizing the responses showed a systematic change in stimulus-evoked charge with bar angle. Orientations that produced the strongest excitation were orthogonal to those producing the strongest inhibition (Figure 5 –figure supplement 2E-F). The same stimulus evoked significantly weaker inward and outward currents from S1-RGCs in Sdk1^CG/CG^ retinae, consistent with their reduced responses to the expanding spot stimulus (Figure 5 –figure supplement 2C-D, G-I). Integrating these responses across stimulus presentation time and normalizing these responses to the average maximal response in controls showed that excitatory currents retained their preference for bar orientation, but the tuning of inhibitory inputs became less selective (Figure 5 – figure supplement 2G-I). Thus, we conclude that Sdk1 is required for S1-RGCs to develop excitatory and inhibitory synaptic inputs, suggesting that Sdk1 promotes synapse formation between these neurons and their interneuron targets.

#### Sdk1 loss impairs S1-RGC dendritic development

Finally, we asked whether there are structural correlates of the reduced synaptic input to S1-RGCs. One possibility is that Sdk1-null S1-RGCs target inappropriate IPL sublayers and therefore cannot receive input from their interneuron partners. However, comparing the laminar position of S1-RGC dendrites in Sdk1^CG/+^ or Sdk1^CG/CG^ retinal cross-sections showed no obvious difference between controls and nulls (Figure 5G-H). We next asked if Sdk1 loss impacts the lateral anatomy of S1-RGC dendrites. We found that the loss of Sdk1 led S1-RGCs to grow dendritic arbors that were less complex than their control counterparts (Figure 5I). Sdk1-null S1-RGC arbors contained similar numbers of dendritic branches (Figure 5J), but they were approximately half as long on average, which led to fewer intersections across the entire dendritic arbor as assessed by Sholl analysis (Figure 5K-M). These deficits arose with no major change in the overall structure of the IPL, as assessed by staining with a variety of markers that label AC and BC subsets targeting specific sublayers (Figure 5 –figure supplement 3). Moreover, reconstructed ONα-RGCs in the same Sdk1^CG/CG^ retina showed dendritic arbors that were similar to their counterparts in control retinae (Figure 5 –figure supplement 1C-H), indicating that Sdk1-loss selectively impairs the dendritic arborization of S1-RGCs.

## Discussion

Here, we investigated the role of Sdk1 in retinal circuit development. In the first section of this study, we molecularly, anatomically, and functionally defined 5 Sdk1 interneurons (ACs and BCs) and 5 Sdk1-RGCs that target an inner set of IPL lamina. This family of RGCs include two ON-DSGC types and ON-OFF S1-RGCs; the latter shows selectivity for edges located in their receptive field. In the second section, we found that Sdk1 loss caused a significant loss of S1-RGC responsivity with little effect on the other Sdk1-RGC types. By comparing visual responses between S1-RGCs and Onα- RGCs in control and Sdk1 nulls, we show that the loss of Sdk1 impairs S1-RGCs’ responses to both bright and dark spot stimuli, as well as oriented bars. Finally, we show that these deficits arise from a selective loss of excitatory and inhibitory synaptic input on S1-RGCs and correlates with a selective loss of small branches in this neuron’s dendritic arbor. We conclude that Sdk1 is required for the dendritic and synaptic development of a local edge-detecting RGC type.

### A family of Sdk1 circuits

Transcriptomic studies indicate that RGCs can be divided into at least 46 different clusters, which include several clusters that do not correspond to any known RGC and are predicted to be novel. In the process of studying Sdk1, we developed molecular signatures for three of these orphan clusters and characterized them using calcium imaging.

We show a pair of Nr2f2+ RGCs can be distinguished based on their expression of Calb and that exhibit selective responses for stimuli traveling ventrally (Nr2f2+/Calb+; C10, (Tran et al., 2019) or dorsally (Nr2f2+/Calb-; C27, (Tran et al., 2019) over the retina. Both cell types exhibit stronger responses to bars whose speed matches the preferred tuning of ON-DSGCs (200μm/sec) and elaborate medium-field dendritic arbors confined to sublamina 4 (Lilley et al., 2019; Matsumoto et al., 2019; Sanes and Masland, 2015). Moreover, reporter-labelled RGCs in the Pcdh9-Cre line, which marks ventral-motion selective ON-DSGCs (Lilley et al., 2019; Matsumoto et al., 2019), colocalize the ventral-motion selective Nr2f2+/Calb+ RGCs. Finally, we show strong RGC axon labelling in the MTN, a brain region known to be targeted by ON-DSGC axons (Dhande et al., 2013; Martersteck et al., 2017; Oyster and Barlow, 1967; Yonehara et al., 2008, 2009). These results strongly suggest that Nr2f2+ Sdk1-RGCs comprise two subtypes of ON-DSGCs.

S1-RGCs are a high-density, narrow dendritic field neuron, characterized by small (<200μm) diameter dendritic arbors, strong surround suppression, and responses to both bright and dark stimuli. Similar properties have been found in several other RGCs that respond to stimuli falling in their receptive field center but are silenced when the same stimuli fall in their surround. One of these RGCs, called W3B, expresses Sdk2 and grows into sublamina 3 just like S1-RGCs, suggesting that the Sdk family plays unique roles in the development of two physically entangled, but functionally distinct local edge-detecting circuits. Four other edge-sensing types are described in a recent atlas of functionally and anatomically defined RGC types. Of these, S1-RGCs most closely resemble type 1 or type 2 high-definition RGCs (HD1 or HD2). Like S1-RGCs, HD1 and HD2 are ON-OFF cells and can respond to edges (Jacoby and Schwartz, 2017). However, S1-RGCs also show sensitivity to edge orientation, which could arise from their orthogonally tuned excitatory and inhibitory inputs. This input arrangement has been observed in recently described horizontally and vertically selective OS-RGCs; however, these neurons show significantly stronger orientation selectivity compared to S1-RGCs. We cautiously speculate that S1-RGCs are a kind of local edge detector with sensitivity to edge orientation. Such a local-orientation selective RGC was reported in a recent calcium imaging survey of RGCs (G14,(Baden et al., 2016), however, a lack of molecular markers in both studies prevent us from drawing a clear correspondence. The molecular markers and genetic access provided by our work offer a way to resolve this issue in future.

### IgSFs and circuit assembly

Several molecular recognition systems direct various steps of retinal circuit assembly, including IPL region selection (Deans et al., 2011; Matsuoka et al., 2011a, 2011b, 2013; Sun et al., 2013) and layer selection (Duan et al., 2014, 2018; Ray et al., 2018; Yamagata and Sanes, 2008; Yamagata et al., 2002). The factors that guide neurons to synapse with targets once they reach appropriate layers are the least characterized, but preliminary work in mammals and invertebrates suggests that IgSF members play a key role in this phenomenon (Carrillo et al., 2015; Cosmanescu et al., 2018; Krishnaswamy et al., 2015; Peng et al., 2017; Sanes and Zipursky, 2020; Tan et al., 2015; Yamagata and Sanes, 2018). Our work here adds to these findings.

We previously showed that an IgSF called Sdk2 enriches connections between VG3-ACs and W3B-RGCs, permitting this RGC to sense object motion (Fosque et al., 2015; Krishnaswamy et al., 2015; Lee et al., 2014). In a separate study, we showed that a different IgSF, Contactin 5 (Cntn5), was required for ON-OFF direction selective RGCs to elaborate dendrites within sublamina 4 and collect input from ACs and BCs (Peng et al., 2017). Too few IgSFs have been studied in the context of mammalian circuit assembly to know whether the differing roles for Sdk2 and Cntn5 represent distinct roles for each IgSF or whether they represent two ends of a continuum in which IgSFs direct small-scale RGC-AC/BC interactions that lead to specific synapses. Here, we show that the closest common relative of Sdk2 and Cntn5, Sdk1, has roles in both phenomena as Sdk1 loss causes selective deficits in both dendritic branching and appearance of functional synapses on S1-RGCs. Interestingly, we did not observe significant phenotypes in RGCs that express both Sdk1 and Sdk2, suggesting that Sdk2 compensates for Sdk1 loss. Taken together, these results point to a common role for IgSFs in promoting or stabilizing the small intralaminar neurites that bear nascent synapses between synaptic partners. Indeed, Sdks localize to synapses through an interaction with the MAGI family of PDZ-scaffolding proteins and can alter dendritic shape if expressed ectopically in early retinal development (Yamagata and Sanes, 2008, 2010). The genetic and molecular definition of Sdk1-RGCs provided by our studies, paired with time-lapse imaging in explants, might offer a way to directly test this idea.

## Acknowledgements

We thank K. Yonehara for gifting us fixed retinal tissue from PCDH9-Cre mice. We thank Drs J. Sanes, E. Cooper, S. Trenholm, K. Yonehara, E. Cook, and E. Feinberg for helpful comments and suggestions on our manuscript.

## Contributions

AK and PLR conceived the study, performed experiments, and analyzed data. AGRO performed gene expression analysis, histological studies, and confocal imaging. CT performed stereotaxic surgeries, histological studies, confocal imaging, and image analysis. AK and PLR wrote the paper.

## Methods

### Animals

Animals were used in accordance with the rules and regulations established by the Canadian Council on Animal Care and protocols were approved by the Animal Care Committee at McGill University. Male and female Sdk1-CreGFP (Sdk1^CG^), Sdk1-CreER (Sdk1^CE^), and Sdk2-CreER mice aged 35-100 days old were used in this study. Details about the generation of these lines can be found in previous studies (Krishnaswamy et al., 2015; Yamagata and Sanes, 2018). Ai27-ChR2-tdTomato mice were obtained from the Jackson Laboratory (AID27, Jackson Labs, RRID: IMSR_JAX:012567) and crossed with Sdk1CreER mice for some anatomical experiments. Vglut2-Cre mice were obtained from the Jackson Laboratory (Jackson Labs, RRID: IMSR_JAX 016963) and crossed with Cre-dependent GCaMP6f lines (AI95D, Jackson Labs, RRID: IMSR_JAX 024105).

### Viruses

AAVrg CAG-flex-GCaMP6f, AAV9 CAG-flex-GCaMP6f, and AAV9 ef1a-flex-Tdtomato viral vectors were purchased from the Canadian Neurophotonics Platform Viral Vector Core Facility (RRID:SCR_016477) and AAVrg-flex-Tdtomato was purchased from Addgene (viral prep: 28306-AAVrg). Retrogradely infecting AAVs were used for brain injections while AAV9 was used for intravitreal injections.

### Injections

AAVs were injected either intraocularly or intracranially to label Sdk1-RGCs. For intracranial injections, mice were anaesthetized using isoflurane (2.5% in O_2_), given a combination of subcutaneous carprofen and local bupivacaine/lidocaine mix for analgesia, transferred to a stereotaxic apparatus, and a small craniotomy (<1mm) made in the appropriate location on the skull using a dental drill. Next, a Neuros syringe (65460-03, Hamilton, Reno, NV) filled with virus was lowered into either the LGN (2.15mm posterior from bregma, 2.27mm lateral from the midline and 2.75 mm below the pia) or SC (3.85mm posterior from bregma, 0.75mm lateral from the midline and 1.5 mm below the pia) using a stereotaxic manipulator. A microsyringe pump (UMP3-4, World precision Instrument, Sarasota, FL) was used to infuse 400nL of virus (15nL/s) bilaterally in dLGN or SC and the bolus allowed to equilibrate for 8 min before removing the needle. For intraocular injections, mice were anaesthetized as above, given carprofen as analgesic and 1μl of virus injected via an incision posterior to the eye’s ora serrata using a bevelled Hamilton syringe (7803-05, 7634-01, Hamilton). Mice were given at least two weeks to recover before experimental use.

### Tamoxifen

Tamoxifen (TMX) (Milllipore Sigma, T5648) was dissolved in anhydrous ethanol at 200 mg/mL, diluted in sunflower oil to 10mg/mL, sonicated at 40°c until dissolved and stored at −20°C. Prior to injection, TMX aliquots were heated to 37°c and delivered intraperitoneally at ∼1g/50g body weight to Sdk1^CE^xAi27D mice. The dose was repeated twice over 2 days and mice were used between 2 to 4 weeks following treatment.

### Histology

Mice were euthanized by isofluorane overdose and transcardially perfused with chilled PBS followed by 4% (w/v) paraformaldehyde (PFA) in PBS and enucleated. Eye cups were fixed for an additional 45mins and brains fixed for an additional 2-5hrs in chilled 4% (w/v) PFA. For whole mount stains, tissue was incubated in primary antibodies for 7 days at 4°C and incubated in secondary antibodies overnight at 4°C following a series of washes. Stains following two-photon calcium imaging were performed similarly but included lectin to stain blood vessels for registration. Identical procedures were used to stain thick brain sections obtained cutting 100μm thick slices from tissue embedded in 2.5% low EEO agarose (Millipore Sigma, A0576) using a compresstome (VF-200-0Z, Precisionary Instruments, Greenville, NC). For cross-sections, post-fixed retinae were sunk in 40% (w/v) sucrose/PBS, transferred to embedding agent (OCT, Tissue Plus™, Fischer Scientific), flash frozen in 2-methylbutane at −45°C and then sectioned onto slides at 30μm thickness on a cryostat. For immunostaining, slides were first washed in PBS, blocked with blocking solution (4% normal donkey serum, 0.4% Triton-X-100 in PBS) for 2h and incubated overnight with Primary antibodies at 4°C. Tissue was then washed with PBS and incubated in secondary antibodies for 2h at room temperature prior to the final wash and tissue mounting.

### Antibodies and blood vessel stains

Antibodies used were as followed: rabbit anti-DsRed (1:1000, Clontech Laboratories, RRID: AB_10013483); chicken anti-GFP (1:1000, Abcam, Cambridge, UK, RRID: AB_300798); mouse anti-Nr2f2 (1:1000, Abcam, RRID: AB_742211); mouse anti-Brn3c (1:250, Santa Cruz Biotechnology, Dallas, TX, RRID: AB_2167543); mouse anti-Nefh (SMI-32, 1:1000, Biolegend, San Diego, CA, RRID: AB_2314912); rabbit anti-Calbindin (1:10 000, Swant, Marly, CH, RRID: AB_2314070); goat anti-Osteopontin (1:1000, R&D Systems, Minneapolis, MN, RRID: AB_2194992); goat anti-VAChT (1:1000, Millipore Sigma, Darmstadt, DE, RRID: AB_2630394); rabbit anti-Calretinin (1:2000, Millipore Sigma, RRID:AB_94259); guinea pig anti-vGlut3 (1:2000, Millipore Sigma, RRID: AB_2819014); guinea pig anti-Rbpms (1:100, Phosphosolutions, Aurora, CO, RRID: AB_2492226); mouse anti-Chx10 (1:300, Santa Cruz Biotechnology, RRID:AB_2216006); mouse anti-Ap2-α (1:100, clone 3b5 from Developmental Studies Hybridoma Bank, Iowa City, IA). Secondary antibodies were conjugated to Alexa fluor 405 (Abcam, RRID: AB_2715515), Alexa fluor 488 (Cedarlane, Ontario, CA, RRID: AB_2340375), FITC (Millipore Sigma, RRID: AB_92588), Cy3 (Millipore Sigma, RRID: AB_92588 or Jackson ImmunoResearch, West Grove, PA, RRID: AB_2340460) or Alexa Fluor 647 (Millipore Sigma, RRID: AB_2687879). Isolectin (Fisher Scientific, Waltham, MA, RRID: SCR_014365) was incubated along with the secondary antibodies when applicable.

### Confocal imaging and analysis

Images of stained tissue were acquired on an Olympus FV1000 confocal laser-scanning microscope or a Zeiss LSM-710 inverted confocal microscope (Advanced BioImaging Facility, McGill University) at a resolution of 512×512 or 1024×1024 pixels with a step sizes ranging from 0.45μm to 8μm. Following image acquisition, minor processing of images including landmark correspondence transforms to match two-photon and stained retinal fields as well as stitching of large tilescans was performed using ImageJ.

For density recovery profile analysis (DRP), RGC soma centers were selected manually on ImageJ and their x-y coordinates used to generate DRPs using code modelled after (Rodieck, 1991). The resulting DRPs were averaged across a given condition and fitted with a sigmoid curve for visual clarity. The mean cell density obtained from this analysis was used to assess the proportion of cells labelled by Sdk1 among BCs, ACs and RGCs.

We analyzed dendritic morphology as described previously (Krishnaswamy et al., 2015). Briefly, images of single S1-RGCs and Onα RGCs taken from P30-60 retinae were traced manually through the z-stack using simple neurite tracer (ImageJ). Path ROIs describing neuronal processes were converted to stacks and analyzed for morphological properties using the Trees toolbox on MatLab (Cuntz et al., 2010).

To obtain mean IPL projection depth of each Sdk1+ type, we took linescans of pixel intensity across the IPL from images stained with antibodies to VAChT (sublamina 2 and 4) or reporter, normalized these signals to the maximum intensity, and averaged these traces across each condition. IPL depth is expressed as a percent and sublaminae judged by the position of the peak intensities in the VAChT channel. We applied the straighten transform (ImageJ) to correct the VAChT bands on a few curved sections and applied the same transform to reporter channels prior to linescan procedure.

### Calcium imaging

Calcium imaging was performed as previously described (Liu et al., 2018). Briefly, mice were dark adapted for at least 2hrs, euthanized and then retinae rapidly dissected under infrared illumination into oxygenated (95% O_2_; 5% CO_2_) Ames solution (Millipore Sigma, A1420). Next, retinae were mounted onto a filter paper with the RGC layer facing up, placed in a recording chamber, mounted on the stage of a custom built two-photon microscope and perfused with oxygenated Ames solution warmed to 32-34 °C. Responses of GCaMP6f+ RGCs to visual stimuli delivered through the objective were imaged at 920nm and collected at 8Hz. Each image plane (360×72μm) of the movie contained GCaMP fluorescence, SR101 fluorescence, stage coordinates, and visual stimulus synchronization pulses to permit offline analysis. A few microliters of sulphorhodamine 101 (SR101, .2mg/mL, Millipore Sigma, S7635) was added to the recording chamber to label blood vessels and a map of the main blood vessels emanating from the optic disk acquired for post-hoc image registration.

### Posthoc image registration and RGC ROI selection

Following each calcium imaging session, averages of each calcium imaged field were stitched into a single image using custom software written in Matlab. These large images contained a map of the blood vessel pattern surrounding the retinal optic disk and imaged fields (Figure 2 –Figure supplement 1A-B). These images were then scaled into microns, overlayed on images of post-hoc stained retinae and rotated or translated until the blood vessel patterns between the two datasets (Figure 2 –Figure supplement 1A-C). Fine blood vessel morphology was then used as inputs to the landmark correspondences function in Fiji to perform final alignment between these two fields (Figure 2 –figure supplement1D-K). ROIs of stained neurons were used to extract responses from two-photon movies (Figure 4 –figure supplement 1L-O).

Osteopontin distribution which is enriched in the dorsal-temporal quadrant of the retina (Bleckert et al., 2014) permitted alignment of RGC calcium responses to the cardinal axes. For a subset of our studies, we were able to re-stain retinae with two additional markers, register these restains as above, and together with the trace similarity score (see below) categorize Sdk1-RGCs.

### Calcium imaging analysis

RGC responses corresponding to stimulus synchronization pulses were extracted, aligned, and analyzed using custom software written in MatLab (Simulink). Responses to stimuli were expressed as a z-score where the mean and standard deviation were obtained from an equal duration of the prestimulus baseline on each trial. We computed three higher descriptive statistics: The On-Off index was calculated as described previously (Baden et al., 2016), but briefly:

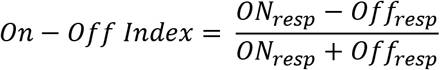

Where ON_resp_ and OFF_resp_ is the mean cell response over a bright and dark full-field stimulus, respectively. ON-OFF indexes from a dataset in which all RGCs express GCaMP6f (Vglut2-Cre::AI27) were plotted alongside Sdk1-RGC ON-OFF indexes after filtering this dataset using a quality index computed as described in Baden et al., 2016. A direction selectivity index was calculated using the circular variance of the cell response for all 8 moving bar direction (Nath and Schwartz, 2016):

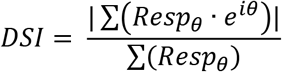

Where Respθ is the maximum cell response for each bar direction. A similar approach was done to compute an orientation selectivity index, where instead of the first complex exponential, the second (*e*^2*i*θ^) is used. A similarity score was used to measure each functional RGC group’s similarity to the group mean. This score is the cosine between the group mean and each individual trace in our dataset. Both traces were euclidean normalized prior to this measurement using the *norm* function in Matlab.

### Electrophysiology

Retinae for electrophysiological recordings were prepared and visualized as described for calcium imaging. For cell-attached recordings, the patch electrodes (4-5MΩ) were filled with Ames solution. For whole cell recordings, patch electrodes (5-7MΩ) were filled with an internal solution containing 112mM Csmethanosulfate, 10mM NaAc, 0.2mM CaCl_2_, 1mM MgCl_2_, 10mM EGTA, 8mM CsCl and 10mM HEPES (pH 7.4). In both cell-attached and whole cell recordings, fluorescein 3000MW dextran (Thermo Scientific, D7156) was added to make the electrode visible under two-photon illumination. For whole cell recordings, internal solution was supplemented with 5mM QX314 Bromide. Excitatory currents and inhibitory currents were isolated by adjusting the holding potential to match reversal potentials for excitation (0mV) and inhibition (E_Cl_ ∼ −60mV). Signals from loose-patch and whole-cell recordings were acquired with a MultiClamp 700B amplifier (Molecular Devices) and digitized at 20khz using custom software written in LabView. For spikes, the Multiclamp was put into I=0 mode and Bessel filter set at 1kHz. For currents, the Multiclamp was put in VC mode and Bessel filter set at 3kHz. Analysis of electrophysiological signals was performed in MATLAB (Simulink) as follows: Briefly, action potentials were detected in loose patch recordings using the *peakfinder* function and binned (50ms) over the entire length of the trial; firing rate histograms for each trial were then averaged and subjected to further processing based on each stimulus. Direction and orientation selective indices were computed from mean firing rate histograms as described for calcium imaging data. For whole-cell currents, trials were averaged, peak amplitude measured, and integral were computed using the *trapz* function across each stimulus epoch. For stationary bar stimuli, integrated currents for each bar flash were normalized to average maximum in controls, plotted against bar orientation for each cell, and then peak responses aligned and averaged across all RGCs in heterozygote or knockout retinae.

### Visual Stimuli

A DLP light crafter (Texas Instruments, Dallas, TX) was used to project monochrome (410nm) visual stimuli through a custom lens assembly which steered stimulus patterns into the back of a 20x objective (Euler et al., 2009). All visual stimuli were written in Matlab using the psychophysics toolbox and displayed with a background intensity set to 1×10^4^ R*/rod/s. Custom electronics were made to synchronize the projector LED to the scan retrace of the two-photon microscope.

For calcium imaging experiments, visual stimuli were centered around the microscope field of view. Moving bar stimuli consisted of a bright bar moving along its long axis in one of eight directions. The bar was 1200μm wide, 3200μm long moving at either 960μm/sec (fast) or 240μm/sec (slow). For electrophysiological experiments, the cell receptive field center was identified using a grid of flashing spots and a small user-controlled probe and the location with the highest response assigned as the center for all subsequent stimuli. Moving bars were thinned for electrophysiological experiments to 200μm; length and speed were the same as calcium imaging studies. Expanding spot stimuli began as a small circle that flashed at the receptive field center and grew after each ON-OFF flash. Stationary, rotating bars had the same dimensions as those for the moving bar but flashed and rotated through 9 orientations. All stimuli were preceded by a gray background whose duration equaled the stimulus duration in each trial.

### RNAseq data analysis

RNA sequencing data was taken from the publicly available Broad institute Single Cell Portal for three previous studies (Shekhar et al., 2016; Tran et al., 2019; Yan et al., 2020). To identify type-specific marker for Sdk1-RGCs we first compared expression of markers between these neurons and all other RGCs to find a differentially expressed subset. Next, we analyzed this subset for the presence 5-gene combinations that could segregate each Sdk1 type; Nr2f2 was identified from Rheaume et al., 2018. Sequencing data plots were generated with the ggplot plot package in R.

### Statistics

No statistical method was used to predetermine sample size. Statistical comparisons between Sdk1-knockout and Sdk1-heterozygote electrophysiological, morphological, and calcium imaging data were performed using the *anova1* function in Matlab (Simulink) followed by the *multcompare* function for pairwise testing.

---

**Figure 1 –figure supplement 1.**
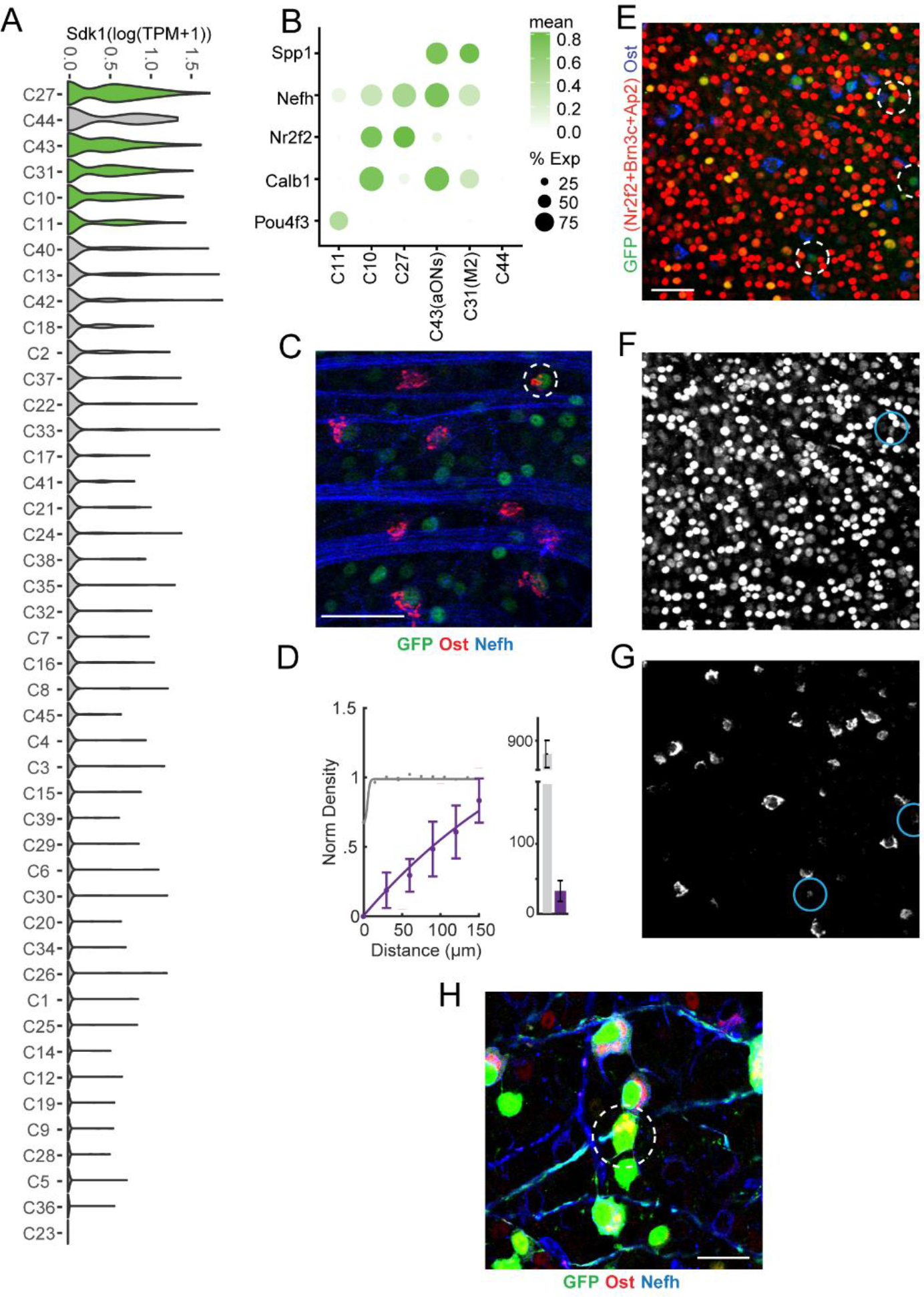
Molecular taxonomy of Sdk1-RGCs. **A.** Violin plots showing expression of Sdk1 extracted from a recently published RNAseq atlas of mouse RGC (Tran et al., 2019). Violins are ordered according to mean Sdk1 expression. Colored violins indicate the RGCs in this study. **B.** Dot plot showing mean expression of marker genes and fraction of expressing cells for Sdk1-RGCs generated from data in Tran et al., 2019. **C.** GFP-stained sample Sdk1^CG^ retinal wholemounts co-stained with antibodies to osteopontin (Ost) and neurofilament heavy chain (Nefh). Dotted circle highlights a single Ost+/Nefh− RGC. Scale = 50μm. **D.** Normalized density recovery profiles and average density of co-labelled RGCs measured from experiments like those shown in C. (n= 13 fields from 6 animals) **E-G.** Image of a Sdk1^CG^ retinal wholemount stained using a cocktail of antibodies against amacrine cells (AP2α), Brn3c, Nr2f2, Ost, and GFP (E), image of the marker cocktail alone (F), and Ost alone with dim marker-labelled cells encircled (G). Greater than 99% of GFP+ cells express the markers shown in E (6 fields from 3 animals). Scale = 50μm. **H.** Wholemount image showing an Ost+/Nefh− RGC amidst a field of other reporter labelled neurons in Sdk1^CG/+^ retinal wholemounts. Scale =25μm.

**Figure 1 –figure supplement 2.**
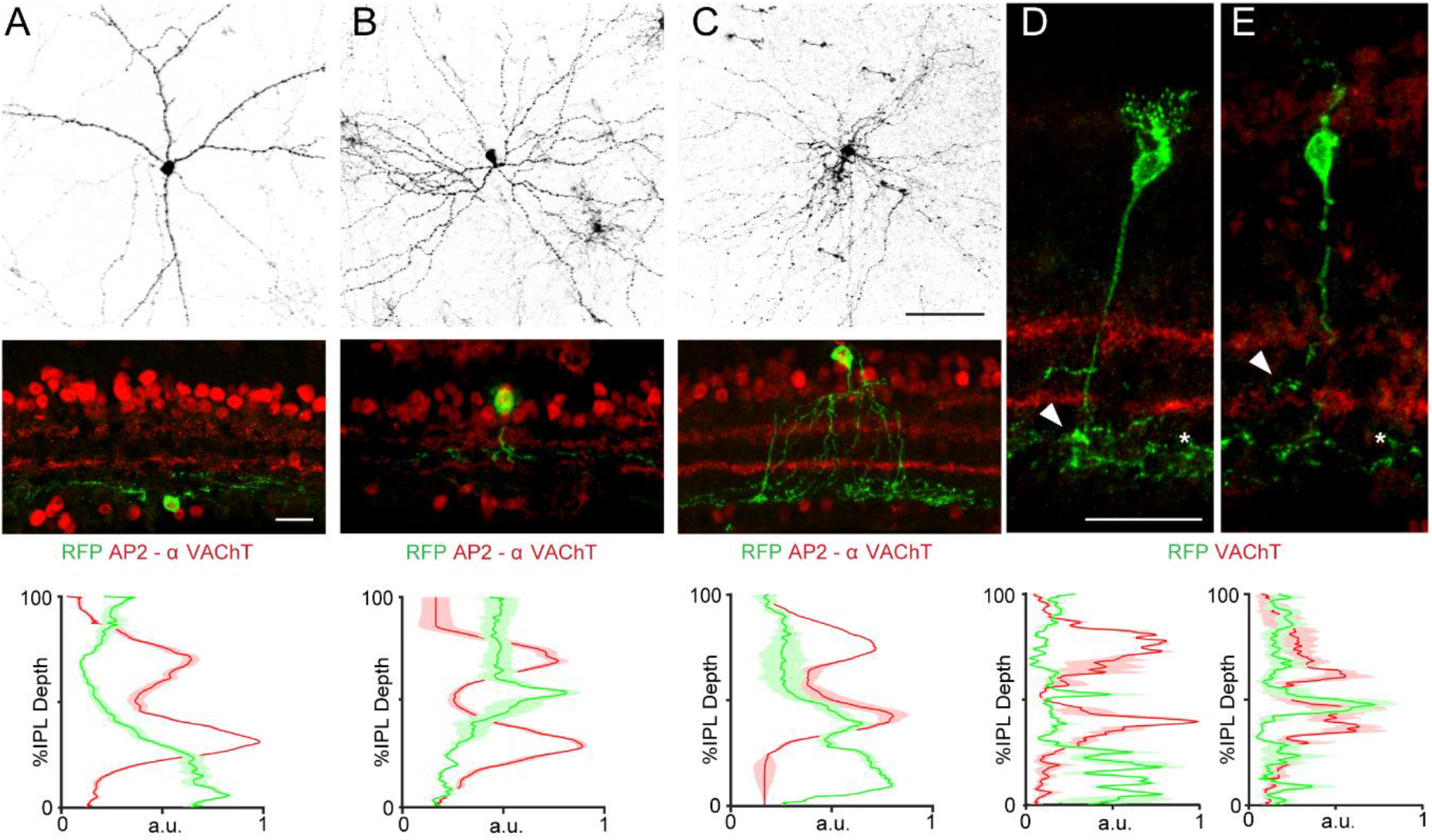
Sdk1 labels 5 kinds of interneurons. **A-C.** Images of Sdk1 amacrine cells in retinal wholemounts (top, Scale = 50μm) and crossections (middle, Scale = 25μm) and corresponding IPL linescans (bottom) showing two widefield types (A-B) and an A17-like waterfall type (C). **D-E.** Images of bipolar cells in retinal crossections taken from Sdk1^CE/+^ mice crossed to reporters whose morphology matches rod bipolar cells (D) or Type7 bipolar cells (E). Arrow heads show BC axon arborization and asterisks denote processes from sublamina-5 bound RGCs and ACs whose somas are not included in this tissue section. Scale = 25μm. (n= 3-6 fields from 2-3 animals for each experiment)

**Figure 2 –figure supplement 1.**
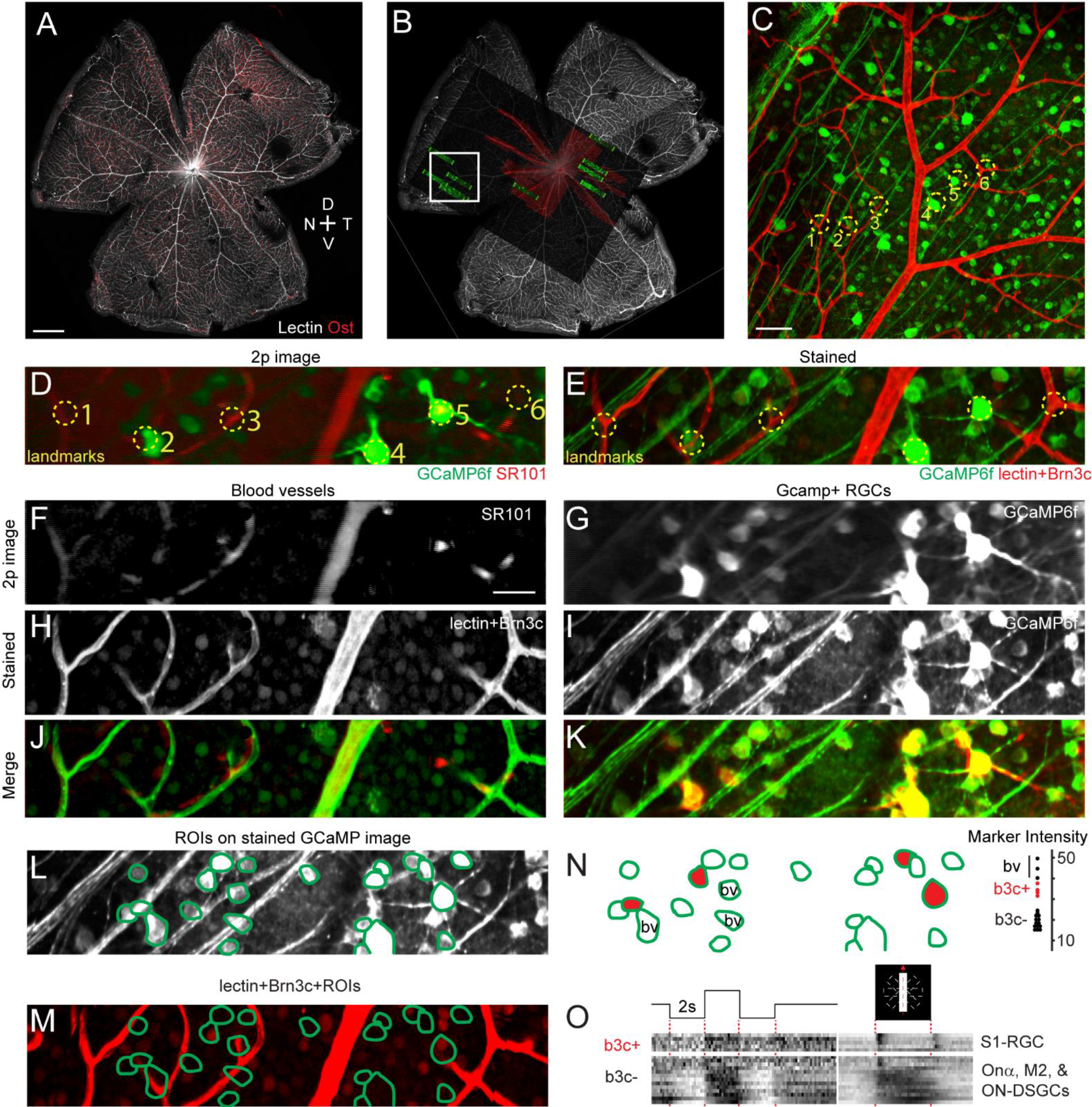
Registration of stained and two-photon imaged retinal fields. **A-C.** Low power wholemount images of retinae stained with lectin and Ost (A) with a superimposed map of calcium imaged fields from two-photon (2p) imaging (B) and a magnified image (C) of the boxed region in B. Lectin staining helps align the two datasets and Ost staining provides retinal orientation (Scale = 500μm) **D-E.** Two-photon field containing the marked neurons in C (D) and confocal image of the same field following staining (E). **F-K.** 2p and confocal images of sulphorhodamine (SR101) or lectin-stained blood vessels and Brn3c (F, H) and GCaMP6f+ RGCs (G, I) shown in D and E and a merge of each pair of images (J, K) showing overlap. **L-M.** Example ROIs drawn on the stained RGC field (L) were used to extract Brn3c marker intensity from a separate channel (M). **N**. The same ROI contours from J with 4 RGCs showing high Brn3c expression (red) and blood-vessel labelling (b.v.). Inset shows pixel intensity for each ROI with those containing high signal due to Brn3c (b3c) or blood vessel labelling indicated. **O.** Mean RGC responses corresponding to the ROIs shown in L to a full-field flash and a bar moving in 8 different directions. High-expressing Brn3c+ RGCs (S1-RGCs) and other Sdk1+ neurons (ONα, M2, ON-DSGCs, and dACs) are indicated. Scale = 50μm.

**Figure 2 –figure supplement 2.**
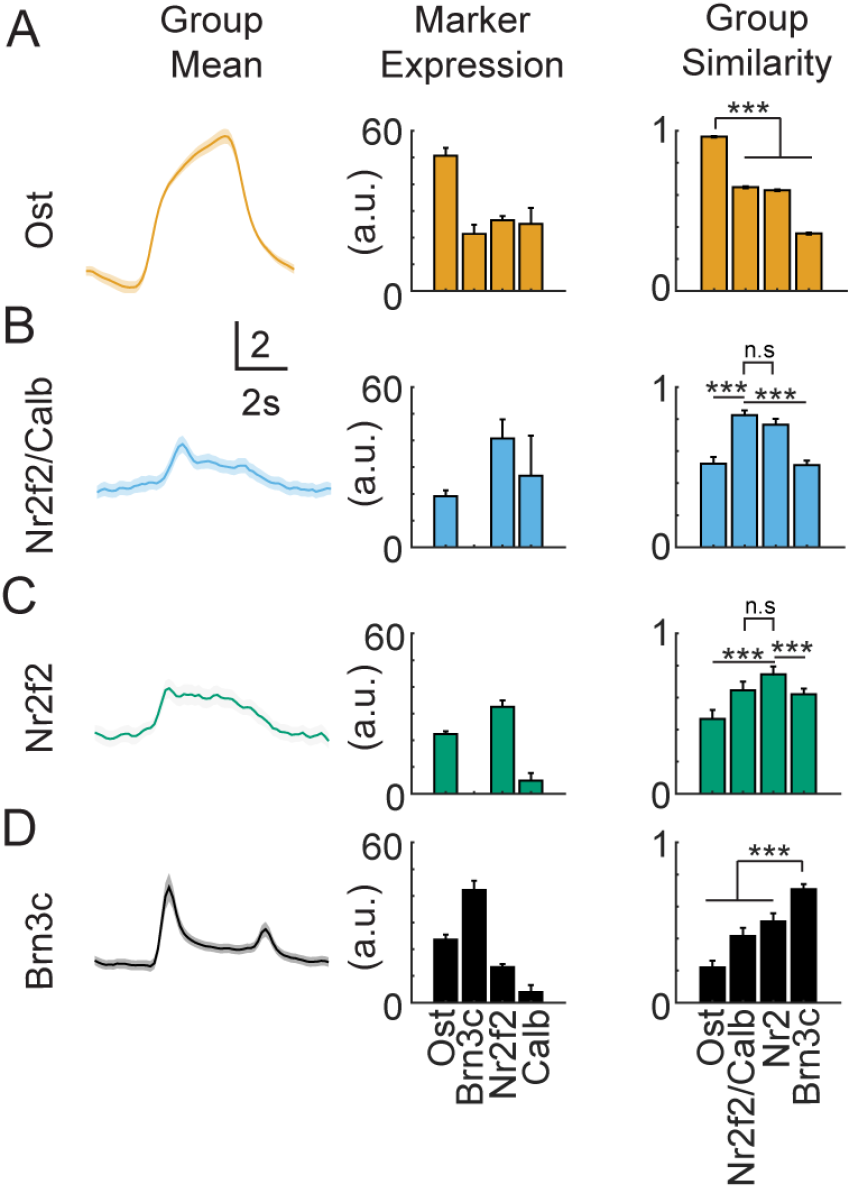
Molecular markers define RGC response clusters. **A-D.** Mean bar response, marker gene expression, and similarity for Ost+ (A), Nr2f2+/Calb+ (B), Nr2f2+/Calb− (C), and Brn3c+ RGC groups (D); vertical scale = z-score of 2. Group similarity bar graphs show the mean of the cosine similarity computed for traces in the Ost (yellow), Nr2f2+/Calb+ (blue), Nr2f2+/Calb− (green), and Brn3c+ groups to the group mean listed on the x axis. Traces are most similar to their group’s mean response versus the mean response for other groups (n values are the same as reported in Figure 2; Significance: *** = p<0.001).

**Figure 3 –figure supplement 1.**
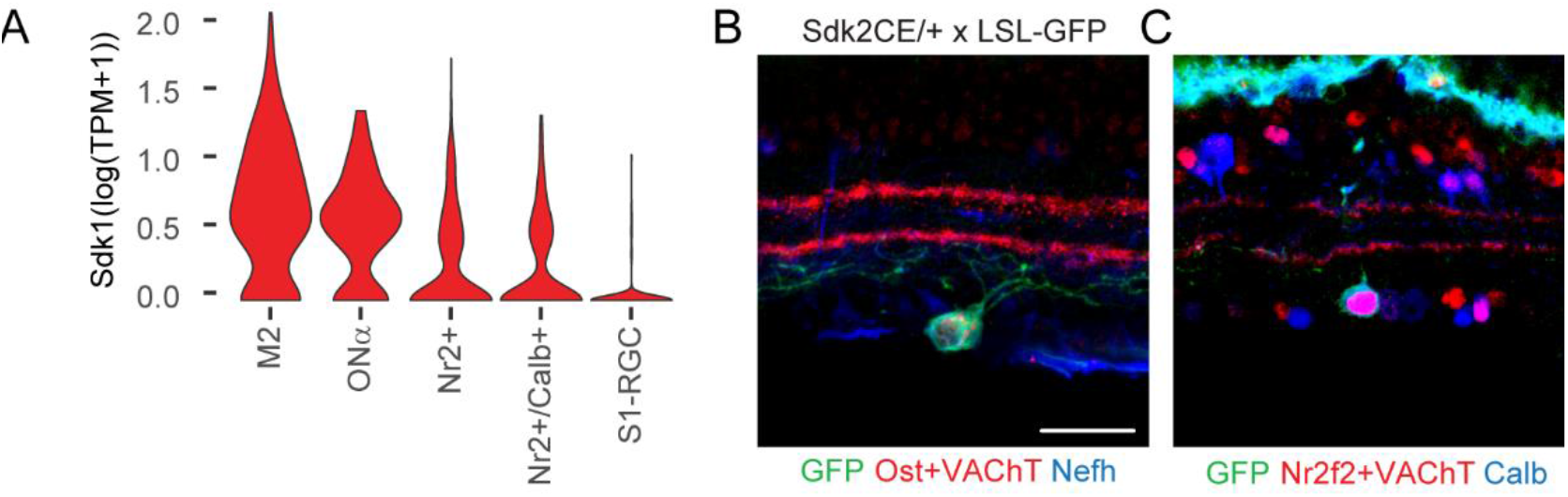
Expression of Sdk2 in Sdk1-RGCs. **A.** Violin plots showing expression of Sdk2 extracted from a recently published RNAseq atlas of mouse RGC (Tran et al., 2019) for the Sdk1-RGCs in this study. Sdk2 expression is nearly absent in S1-RGCs. **B-C.** Image of an Ost+/Nefh+ Onα-RGC and an Nr2f2+/Calb+ RGC in retinal cross-sections taken from Sdk2^CE/+^ mice crossed with Cre-dependent reporters. Scale = 30μm.

**Figure 5 –figure supplement 1.**
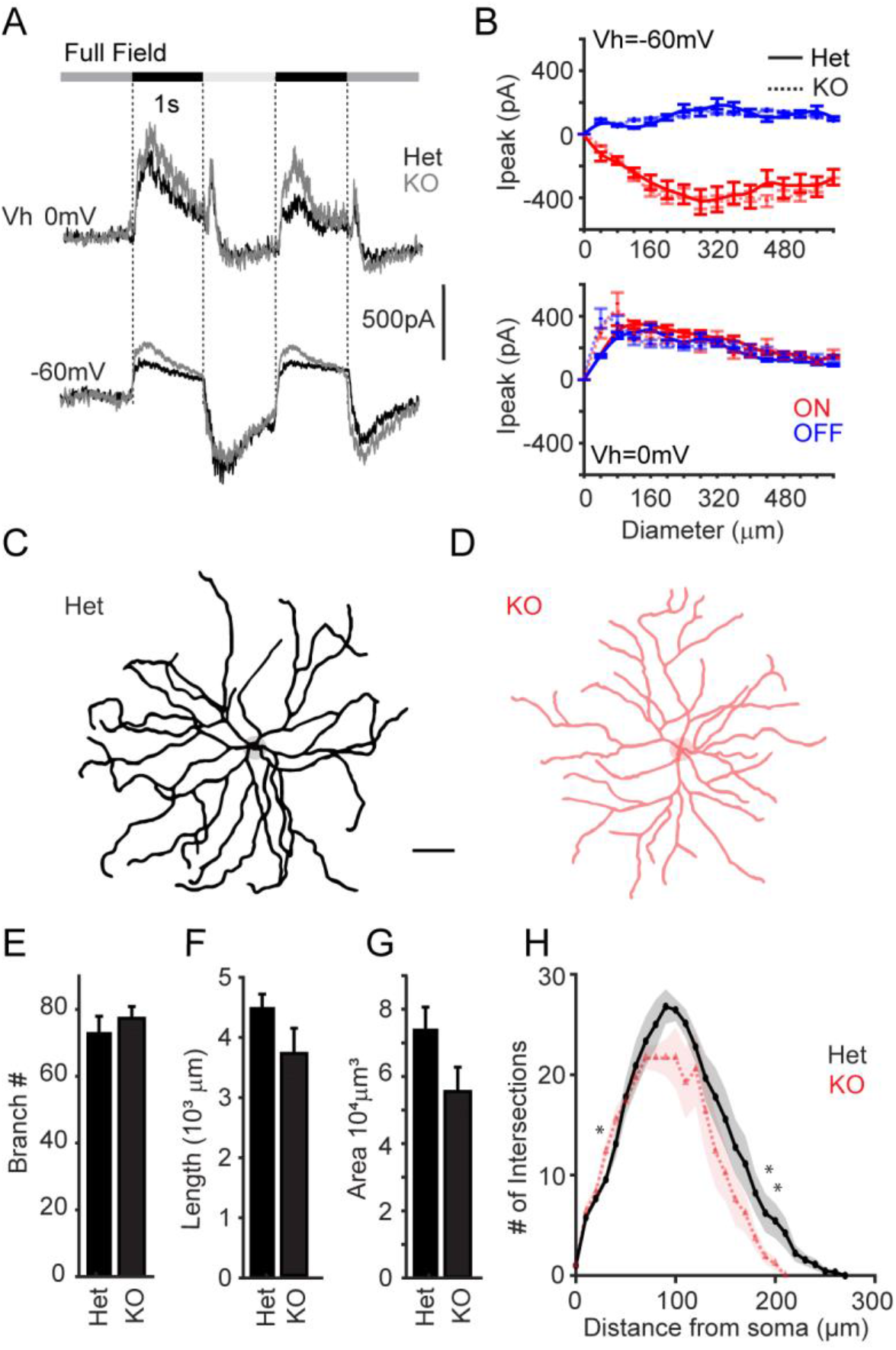
Sdk1+ Onα-RGC synaptic inputs and dendritic arbors are unaffected by Sdk1 loss. **A.** Whole cell recordings from ONα-RGCs held at potentials to isolate excitatory (∼-60mV) and inhibitory (∼0mV) synaptic currents to a full field flash in Sdk1^CG/+^ (Het) or Sdk1^CG/CG^ retinae (KO). **B.** Average peak excitatory (∼-60mV, top) and inhibitory (∼0mV, bottom) currents versus expanding bright (ON) or dark (OFF) spot diameter measured from Het or KO ONα-RGCs presented with expanding spot stimuli. (n=4 for Sdk1-Het, n=5 for Sdk1-KO) **C-D.** Skeletonized ONα-RGC dendrites from control (black) and Sdk1 null (red) retinae. (Scale bar = 50μm). **E-G.** Average branch number (E), branch Length (F), and dendritic area (G) measured from control and Sdk1 null S1-RGC dendritic arbors (L). **H.** Sholl analysis of dendritic arbors measured from Het and KO ONα-RGCs. (n = 9 Sdk1-Het and n=7 Sdk1-KO; * = p < 0.05; ** = p < 0.01)

**Figure 5 –figure supplement 2.**
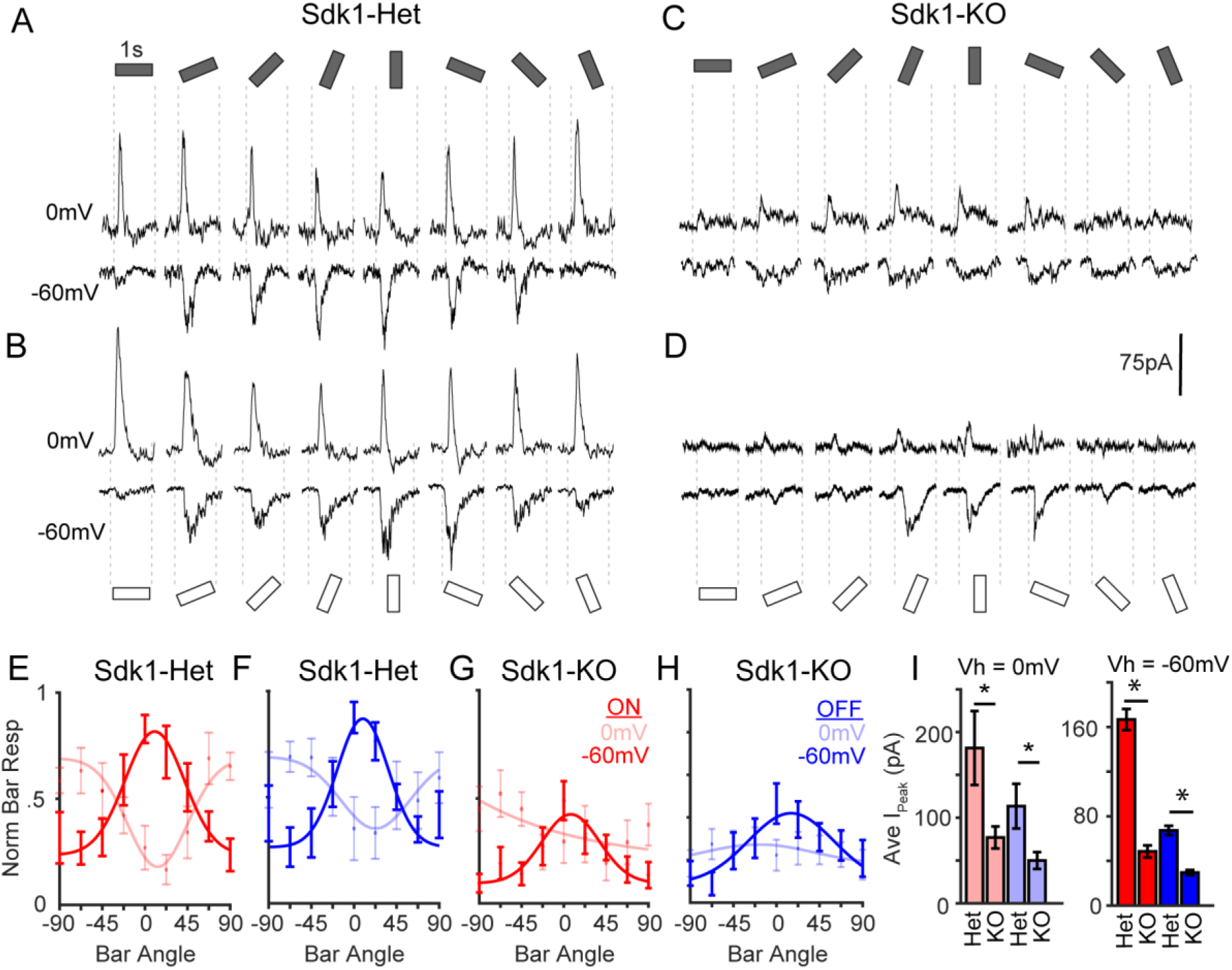
Synaptic currents evoked by oriented bars in control and Sdk1-null S1-RGCs. **A-D.** Sample whole cell currents measured from S1-RGCs held at potentials to isolate excitatory (∼-60mV) and inhibitory (∼0mV) synaptic inputs to a dark (blue) or bright (red) bar that rotated through 8 different angles in Sdk1^CG/+^ (Het) or Sdk1^CG/CG^ retinae (KO). **E-H.** Normalized responses versus the orientation of bright (E, G) or dark (F, H) bars presented to S1-RGCs in control (E-F) or KO (G-H) retinae. Y-values equal the stimulus-evoked charge integrated across the entire bar presentation time and normalized to the average maximum in controls. **I.** Average peak current (I_peak_) evoked by bright (blue) or dark (red) bars for Het or KO S1-RGCs in the experiments shown in A-D. (n=8 for Sdk1-Het, n=9 for Sdk1-KO; * = p < 0.05)

**Figure 5 –figure supplement 3.**
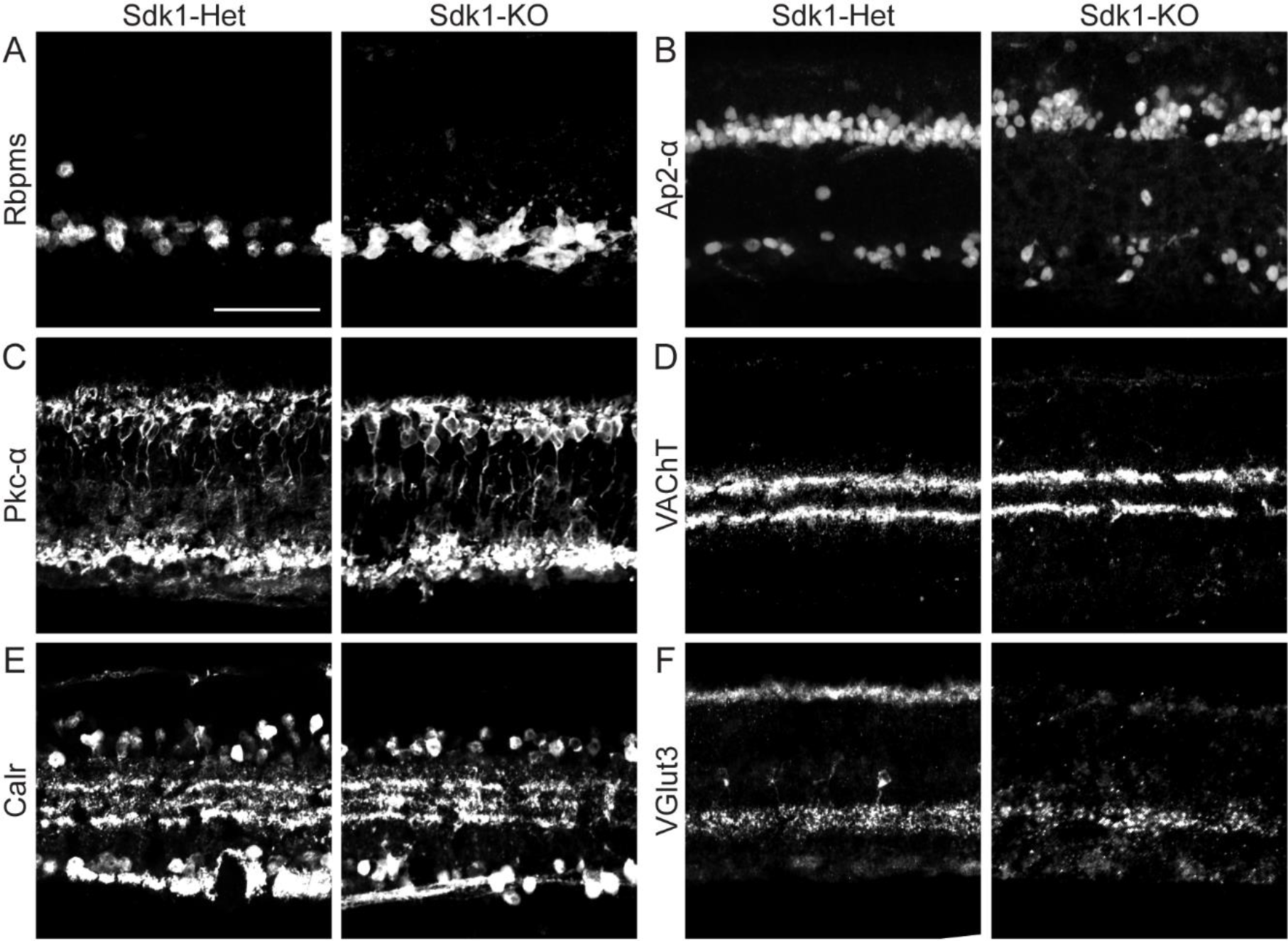
IPL lamination in Sdk1 null retinae resembles controls. Sample retinal cross sections stained with Rbpms (A), Ap2α (B), Pkc-a (C), VAChT (D), Calretinin (E) and VGlut3 (F) taken from Sdk1^CG/+^ (Het) or Sdk1^CG/CG^ retinae (KO) (Scale = 50μm).

